# Broad heterologous protection against Influenza A viruses by an adjuvant-free modular mucosal T-cell vaccine platform

**DOI:** 10.64898/2026.03.29.715080

**Authors:** Rajesh T. Yadav, Mansi Sharma, Santhosh K. Nagaraj, Rohan Narayan, Abinaya Kaliappan, Uma Shanmugasundaram, Rahul Chavan, Akhila B. Rai, Rashmi Rai, TSK Prasad, Suryaprakash Sambhara, Rachit Agarwal, Shashank Tripathi

## Abstract

Rapid antigenic evolution of Influenza A viruses (IAVs) enables their escape from strain-specific vaccine immunity and underscores the need for broadly protective strategies. Here, we describe a modular, adjuvant-free mucosal vaccine platform that elicits potent and cross-protective T cell immunity. The approach uses overlapping CD4+ and CD8+ epitope-dense regions from the consensus IAV M1 and NP proteins, identified through computational and functional screening. These peptides are delivered using polylactic-co-glycolic acid (PLGA) microparticles, engineered for selective uptake by antigen-presenting cells and enable sustained, pH-responsive antigen release. This design enhances antigen processing and MHC cross-presentation, functionally substituting for a conventional adjuvant. This formulation drives robust activation of primed human as well as murine CD4+ and CD8+ T cells and confers broad protection against homologous (H1N1, H3N2) as well as heterologous (H5N1) IAV strains in immunized mice. Overall, this adjuvant-free dose-sparing platform establishes an adaptable framework for next-generation broadly-protective vaccines against rapidly-evolving viruses.

**Graphical abstract:** 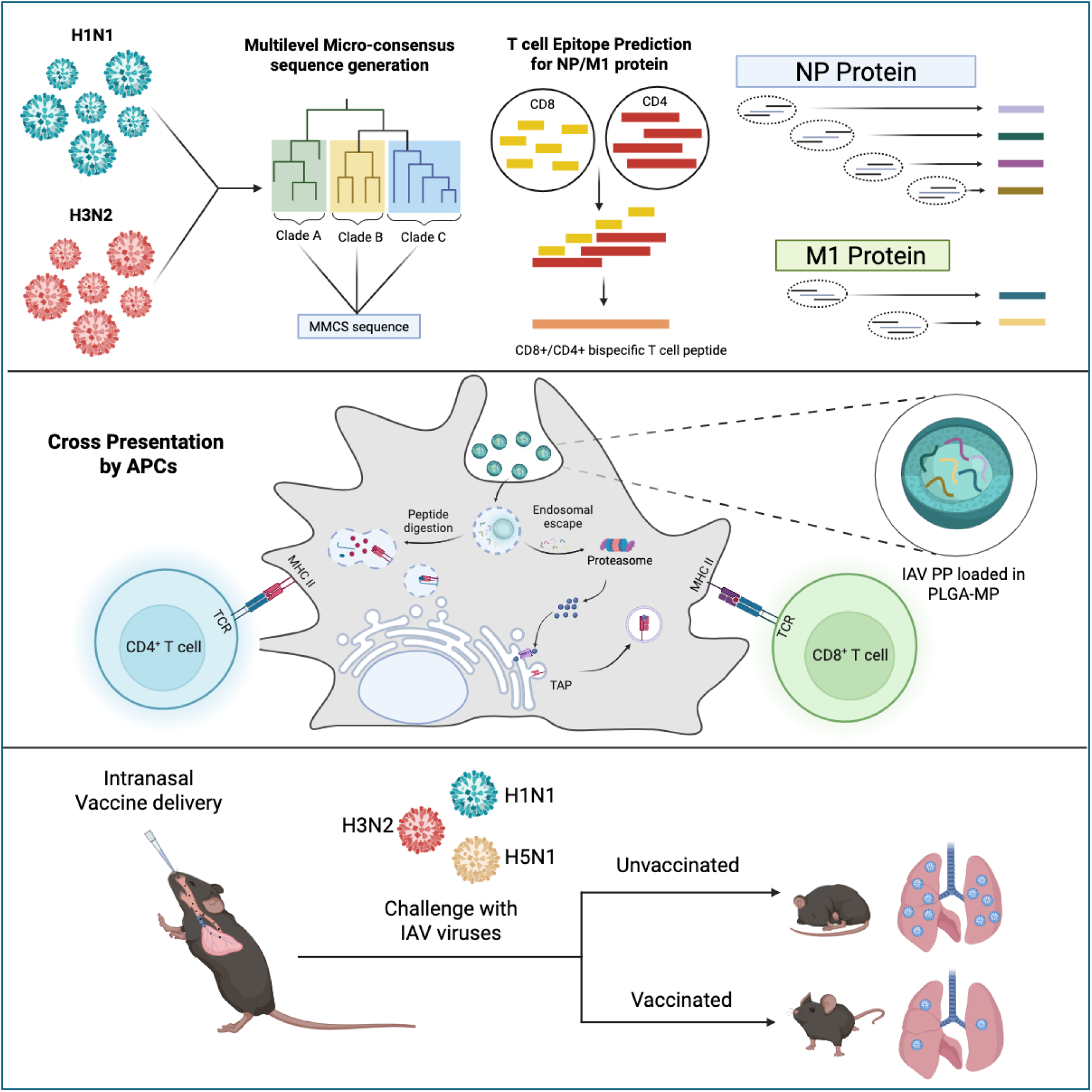

## INTRODUCTION

Influenza A virus (IAV) remains a major global public health threat, causing widespread respiratory infections and significant morbidity and mortality each year, particularly among vulnerable populations (*1*). The World Health Organization estimates that seasonal influenza infects about 1 billion people annually, leading to 3-5 million severe cases and 290,000-650,000 deaths (*2*). Rapid antigenic evolution enables the virus to evade immunity, while current vaccines provide only moderate and variable protection (∼40-60%), depending on how closely vaccine strains match circulating viruses (*3*). Existing flu vaccines primarily target the surface glycoproteins hemagglutinin (HA) and neuraminidase (NA); however, this strategy is undermined by frequent antigenic drift and occasional shifts (*4*), necessitating the annual reformulation of vaccines (*5*). Moreover, reliance on egg-based production systems can introduce manufacturing delays and adaptive mutations in the virus, further diminishing vaccine effectiveness (*6*).

To address these limitations, considerable efforts have focused on developing broadly protective vaccines against IAVs, which remains a major global objective. Most approaches have aimed to elicit broadly neutralizing antibodies against conserved regions of HA; however, these strategies remain susceptible to antigenic variation and often require complex antigen design. Comparatively few studies have explored vaccines that harness cellular immunity, particularly peptides encoding CD8+ T cell epitopes as candidate immunogens (*7*). Although T cell epitopes are relatively conserved, their application as vaccine candidates faces key challenges: they are prone to proteolytic degradation, rapidly cleared from circulation, and require potent adjuvants to promote efficient cellular uptake and effective antigen presentation (*8*). Exogenous peptide antigens are typically processed through the MHC class II pathway, which limits their ability to induce the MHC class I dependent cytotoxic T cell responses required for effective antiviral immunity.

Emerging evidence further highlights the importance of coordinated CD4+ and CD8+ T cell responses in antiviral immunity. While CD4+ T cells play a key role in viral clearance, they also contribute independently to protection and are essential for an optimal immune response. Indeed, depletion studies demonstrate that the absence of both subsets severely impairs viral clearance (*9*). Moreover, non-lethal infections can occur, even in the absence of CD8+ T cells, underscoring the crucial role of CD4+ T cells in antiviral defense (*10*). Recent studies further demonstrate that CD4+ T cells are key contributors in mitigating disease severity in flu infection (*11*). These findings underscore the importance of vaccines that can induce coordinated CD4+ and CD8+ T cell responses.

An important but underexplored aspect of T-cell vaccine design is the organization of epitopes within antigenic regions. Current T cell vaccine strategies fail to provide virus protection because they rely on isolated epitopes that offer limited breadth and elicit only a CD8+ T cell response. Viral proteins often contain clusters of overlapping CD4+ and CD8+ epitopes that may facilitate coordinated antigen processing and presentation. We therefore hypothesized that selectively targeting such epitope-dense, conserved viral protein regions, rather than isolated epitopes, enriched in both CD4+ and CD8+ T cell epitopes, could enhance immunogenicity, improve HLA coverage, and enable more effective cross-subtype protection.

In this context, we developed a novel T cell-focused vaccine strategy based on evolutionarily conserved regions of IAV abundant internal proteins, the nucleoprotein (NP) and matrix 1 (M1) proteins derived from H1N1 and H3N2 subtypes. We applied a multi-level micro-consensus (MMCS) approach to generate representative consensus sequences of the internal proteins NP and M1, minimizing subtype bias. Using a bioinformatics pipeline, we predicted immunodominant CD8+ and CD4+ T cell epitopes with broad HLA coverage, identifying 16 conserved regions in NP and M1 consensus sequences that are highly enriched for T cell epitopes. Functional validation led to the selection of six bispecific peptide regions, two from NP and one from M1, for each subtype, each containing overlapping CD4+ and CD8+ epitopes.

To overcome the delivery limitations of peptide vaccines, these peptides were encapsulated in biodegradable polylactic-co-glycolic acid microparticles (PLGA-MPs), enabling sustained release and efficient targeting of antigen-presenting cells (APCs). This system-enhanced peptide uptake facilitated low-pH-dependent intracellular release and promoted cross-presentation via the MHC-I pathway. PLGA-encapsulated peptide pool (PP) induced robust CD4+ and CD8+ T cell responses in human and murine systems. Importantly, adjuvant-free intranasal immunization was safe and conferred protection in mice against homologous H1N1 and H3N2 viruses, as well as against heterologous H5N1 challenge, demonstrating broad protective potential. Overall, this study presents a first-in-class, adjuvant-free, intranasal T cell-based vaccine platform that safely induces broad protective immunity against IAV, representing a promising step toward next-generation influenza vaccines.

## RESULTS

### Generation of an evolutionary micro-consensus of NP and M1 proteins of human H1N1 and H3N2 IAVs

The NP and M1 proteins of IAV are significant virion constituents produced in high copy numbers during infection (*12*). They are relatively conserved compared to surface glycoproteins and are prominent targets of T cell-mediated immunity (*13*). According to our inclusion criteria (Figure 1A), we had 1,574 H1N1 and 1,742 H3N2 NP sequences, along with 694 H1N1 and 443 H3N2 M1 sequences. The global distribution of our selected sequences showed coverage of all inhabited continents, with the highest representation from North America and the lowest from Africa (Supplementary Fig. 1A). To understand the evolutionary diversity of compiled sequences, we generated a neighbor-joining phylogenetic tree using HA protein sequences (Figure 1B). NP and M1 sequences were mapped radially, with the innermost ring indicating the isolation year, the second ring indicating the HA clade classification, and the outer layers indicating the NP and M1 sequences (Figure 1B). This analysis showed temporal and evolutionary coverage across major IAV clades. Next, to generate a single evolutionary consensus for NP/M1 proteins of H1N1/H3N2 subtypes, we used a MMCS method to avoid clade-specific biases among different sequence groups (Supplementary Fig. 1B-E). Our approach adapts the computationally optimized broadly reactive antigen (COBRA) method, previously used for HA nucleotide consensus sequences in vaccine development (*14*). To verify that the final MMCS consensus represents a more accurate evolutionary average of selected NP and M1 protein sequences, we compared it with an overall conventional consensus and clade-specific consensus sequences (Figure 1C-F). The analysis revealed that the MMCS consensus of NP/M1 showed greater homology with most clade-specific consensus sequences than the overall conventional consensus sequences. The final MMCS consensus sequences were then input into the T cell epitope prediction pipeline for downstream analysis.

**Figure 1.**
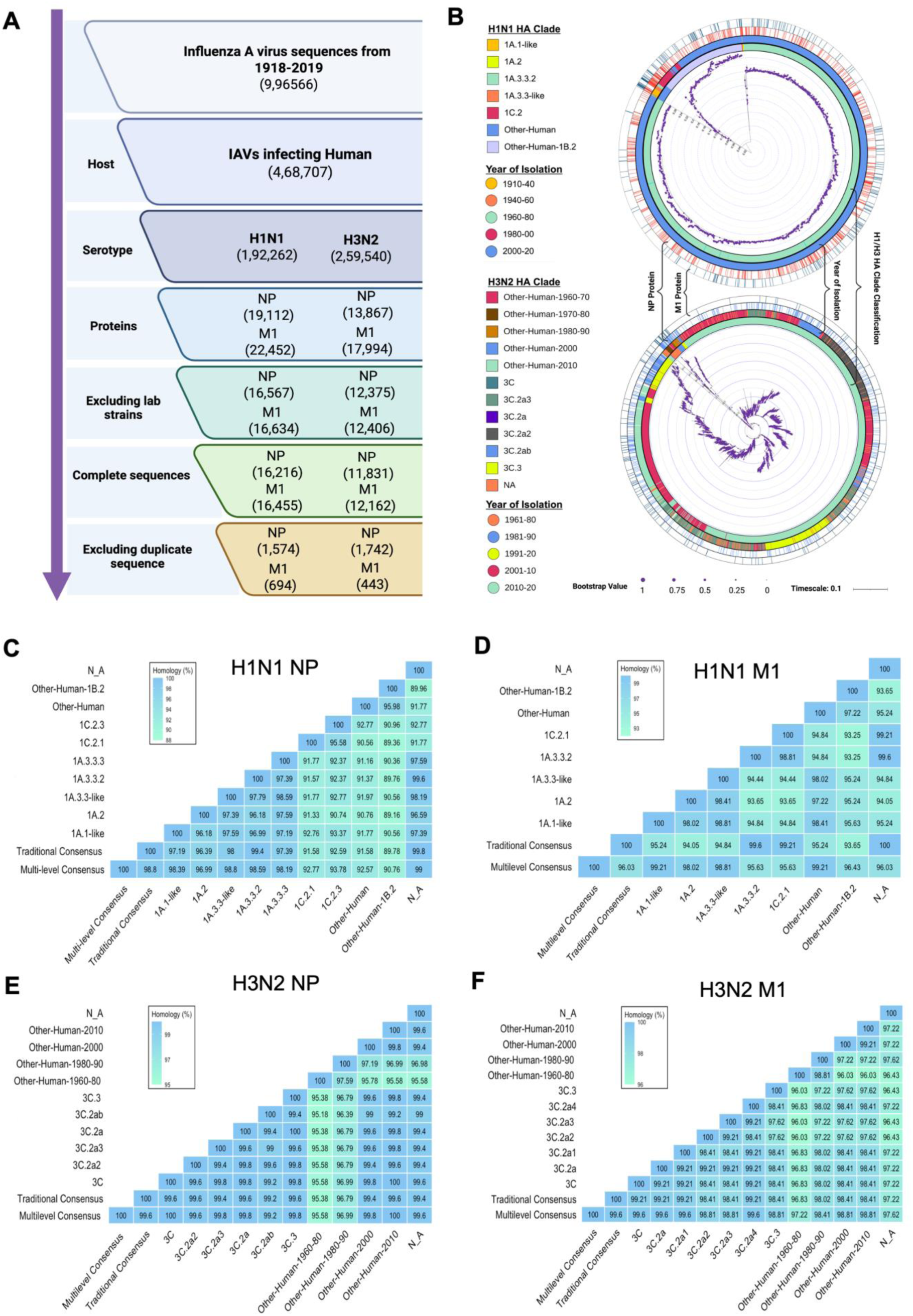
Selection criteria, Evolutionary analysis, and Homology comparison of NP and M1 protein sequences from H1N1 and H3N2 human IAV: **(A)** Data filtering pipeline for selecting representative IAV sequences. The process begins with a million sequences (1918-2019) and narrows to unique, complete human H1N1 and H3N2 NP and M1 protein sequences. **(B)** Circular HA protein phylogenetic tree of H1N1 (top) and H3N2 (bottom) IAV. The inner rings provide metadata layers including year of isolation, HA clade classification, NP, and M1 sequence identity groupings. Heatmaps showing pairwise sequence homology (%) comparisons were generated using traditional consensus with MMCS sequences against known H1N1 **(C, D)** and H3N2 **(E, F)** viral clades for the NP and M1 proteins.

### Prediction and mapping of immunodominant CD8/CD4 bispecific regions in the NP/M1 proteins of diverse IAVs

The T cell immunity-based IAV vaccine development efforts have focused primarily on CD8+ epitopes of viral proteins (*15*). We hypothesized that including CD4+ epitopes, in addition to CD8+ epitopes, could further enhance vaccine efficacy. Additionally, instead of focusing on individual epitopes, viral protein regions saturated with multiple CD8+ and CD4+ epitopes can provide a more robust and broad-spectrum immune response. To test this, we used a rationally designed bioinformatic pipeline to identify such regions on the MMCS NP and M1 protein sequences generated earlier (Figure 2A). Our approach involved the use of three highly cited T cell epitope prediction tools (IEDB I/II, NetMHCpan I/II, and Propred I/II), and the selection of epitopes that were positively predicted by all three (*16–18*). Each tool employs a distinct algorithmic framework, such as ANN-based, matrix-based, and pan-allelic modelling, ensuring that a single computational approach does not skew predictions. A significant challenge in T cell epitope prediction is that MHC alleles exhibit extensive diversity and varying frequencies across different population groups worldwide (*19, 20*). Using a compilation of three tools ensured extensive coverage of both common and rare HLA alleles. In summary, the predicted epitopes provided over 97% coverage of class I and 99% coverage of class II HLA alleles listed in the IEDB reference set (Table S1) (*19, 21*). Initially, around 0.1 million peptides (9-mer for MHC I and 15-mer for MHC II) were screened for HLA binding. A stringent cut-off was applied to the top 2% percentile for MHC I and the top 10% for MHC II, resulting in 4098 predicted CD8+ and 7392 predicted CD4+ T cell epitopes. From these, only peptides predicted as top binders by all three tools were retained, yielding 412 CD8+ and 1,517 CD4+ candidate epitopes. To ensure proper antigen processing, predicted peptides should also exhibit favorable proteasomal cleavage (typically a score≥ 0.5) and efficient TAP transport, with higher scores indicating better ER translocation. For immunogenicity, peptides with positive scores (score > 0) are generally favored. To avoid triggering autoimmunity, peptides are screened against the human proteome, and those that share an identity match of 80-90% or more with human proteins are excluded. Furthermore, we selected only epitopes for which experimental evidence is indicated in IEDB, resulting in 112 epitopes. The selected CD4+ and CD8+ T cell epitopes were then mapped onto the NP and M1 MMCS protein sequence in ascending order of their positions. This led to the selection of 16 CD8/CD4 bispecific epitope-enriched regions (Figure 2B). We also predicted the ability of the selected CD8+ and CD4+ epitopes to be recognized by the MHC of C57BL/6 mice. Interestingly, all 16 CD8/CD4 bispecific regions contained epitopes recognized by the C57BL/6 mouse MHC, suggesting they may be immunogenic in the murine model. We also investigated the potential for cross-reactive immunity of these peptides against other IAV subtypes and found that they provide 100% coverage against most IAV strains across multiple subtypes (Table S2).

**Figure 2.**
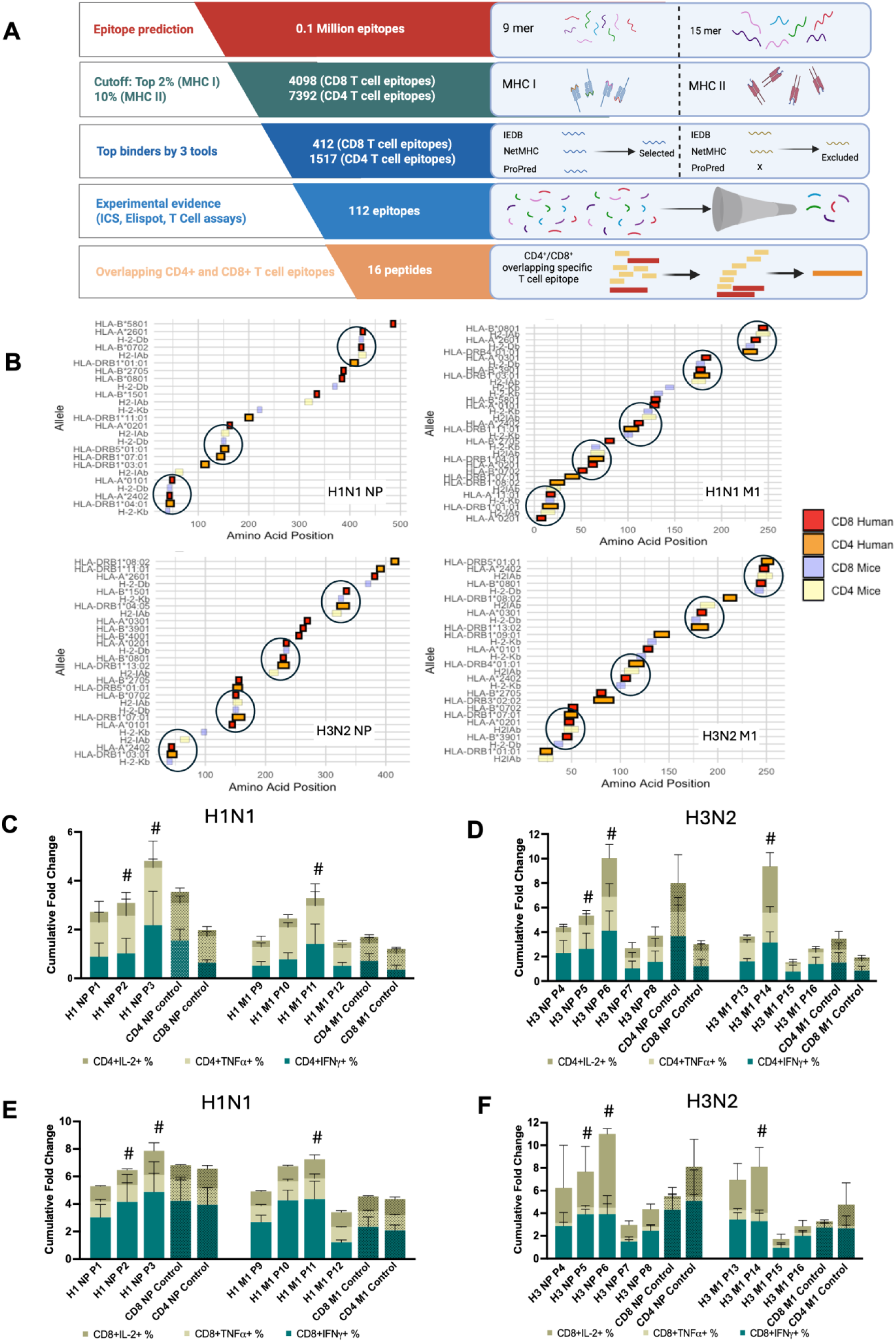
Prediction and Mapping of CD4+ and CD8+ Enriched Bispecific T Cell Epitopes in NP and M1 Proteins of Human IAVs: **(A)** Workflow for identifying immunogenic T cell epitopes using a multilevel filtering approach. **(B)** Positional mapping of selected CD4+ and CD8+ T cell epitopes in the NP and M1 proteins of ma/Cal/09 H1N1 and HK/19 H3N2 viruses. Colored bars indicate epitope across species (human/mouse) and T cell types (CD8/CD4). **(C-F)** Cumulative fold-change in CD4+ (C, D) and CD8+ (E, F) T cell responses (based on IL-2+, TNF-α+, and IFN-γ+ populations) measured by flow cytometry in response to peptide stimulation for H1N1(C, E) and H3N2 (D, F) infected mice splenocytes (n=6-8). Comparisons include individual NP/M1 peptides and control epitopes for both H1N1 (C, D) and H3N2 (E, F). # indicates the T cell peptides selected for our study.

### Functional screening and selection of immunodominant CD8/CD4 bispecific peptides derived from NP and M1 consensus proteins of human H1N1 and H3N2 IAVs

Next, we tested the immunogenicity of the CD8/CD4 bispecific peptides derived from the NP/M1 IAV peptides shortlisted in the previous analysis. We followed the experimental outline depicted in Supplementary Fig. 2A to assess the ability of these peptides to activate both CD8+ and CD4+ T cell responses. Briefly, C57BL/6 mice were infected with 1xMLD_50_ of mouse-adapted A/California/04/2009 H1N1 (ma/Cal/09), A/Hong Kong/2671/2019 H3N2 (HK/19), and A/X-31 H3N2 (X31, a reassortant virus with internal genes from A/Puerto Rico/8/1934, PR8 H1N1 strain). Post 8 days of infection, splenocytes were collected and stimulated with 10 µg of individual peptides, and the expression of IFN-γ, TNF-α, and IL-2 cytokines in activated CD8/CD4 T-lymphocyte subsets was measured by intracellular cytokine staining (ICS) using flow cytometry (Supplementary Fig. 2B-C).

The results showed a significant upregulation of cytokines in splenocytes from infected mice compared with uninfected controls (Figure 2C-E). Notably, all 16 peptides activated CD8+and CD4+ responses comparable to or higher than the control peptides (Figure 2C-E). Finally, P2, P3, and P11 peptides for H1N1 viruses, and P5, P6, and P14 peptides for H3N2 viruses were selected based on their superior potential to activate bispecific CD8/CD4 T cells. The six chosen IAV peptides and the respective CD8/CD4 epitopes they contain are listed in Table S3.

### Encapsulation of IAV peptides in PLGA-MPs promotes controlled release and efficient APCs uptake

A significant challenge for T cell peptide-based vaccines is enzymatic degradation in the biological environment before they are taken up by APCs. PLGA is an FDA-approved, biodegradable, and biocompatible polymer, making it a highly suitable delivery vehicle for clinical applications (*22*). Peptide encapsulation in PLGA-MPs can shield them from proteolytic degradation in the bloodstream or tissues. Considering this, we encapsulated the 6 selected IAV peptides into PLGA-MPs using the double emulsion method (water/oil/water) (*23*). Scanning Electron Microscopy (SEM) imaging and fluorescence microscopy revealed that the PLGA-MPs were spherical, with an average size range of 1-2 μm (Figure 3A-B). Dynamic light scattering (DLS) analysis showed a narrow size distribution with an average diameter of 1450 ± 211.6 nm and a low polydispersity index (0.183), indicating a relatively homogenous MPs with minimal size variability (Figure 3C). This size range of PLGA-MPs is preferred for uptake by APCs, as particles in the micron size range are efficiently internalized through phagocytosis, thereby enhancing intracellular antigen processing and subsequent presentation via MHC molecules (*24*). Efficient peptide loading is critical to minimize immunogen loss and accurately, determine the PLGA-MP dose required for immunization. In our formulation, we achieved an encapsulation efficiency (EE) of 15-20%, corresponding to a loading level (LL) of 8-12.5 µg of IAV PP per mg of PLGA-MPs (Figure 3D), which is consistent with previously reported values (*25*). PLGA-MPs are internalized by APCs via phagocytosis and endocytosis, during which the intravesicular pH progressively drops. To mimic the pH conditions encountered in endocytic and lysosomal compartments, we evaluated the impact of acidic pH on the release kinetics of IAV peptides from PLGA-MPs. We measured in vitro release kinetics of the PLGA-MPs at pH 5 and 7 (Figure 3E). We observed ∼70% peptide release at pH 5 and ∼44.8% at pH 7 through day 4. By day 14, ∼93.5% of peptides were released at pH 5, compared to 61.6% at pH 7. These findings demonstrate that PLGA-MPs provide sustained peptide release, with accelerated release under acidic conditions that simulate the intracellular environment following APC uptake, thereby facilitating efficient intracellular release of the peptide cargo. Next, we evaluated the efficiency of peptide uptake from PLGA-MPs by APCs, using a FITC-labelled peptide (P6) and examining its uptake in mature bone marrow-derived dendritic cells (BMDCs) isolated from naïve C57BL/6 mice. The mature BMDCs were characterized using the surface markers F4-80-, CD11c+, CD80+, CD86+, and MHC-II+ (Supplementary Fig. 3). The results showed that while the free peptide exhibited approximately 80% and 18% uptake for 10 µg and 2.5 µg doses, respectively, corresponding doses of the FITC-tagged peptide encapsulated in PLGA-MPs achieved significantly higher uptake (approximately 95% for 10 µg and 58% for 2.5 µg) (Figure 3F-G). These findings demonstrate that encapsulating the peptides in PLGA-MPs enhances cellular uptake compared to the free peptide formulation.

**Figure 3:**
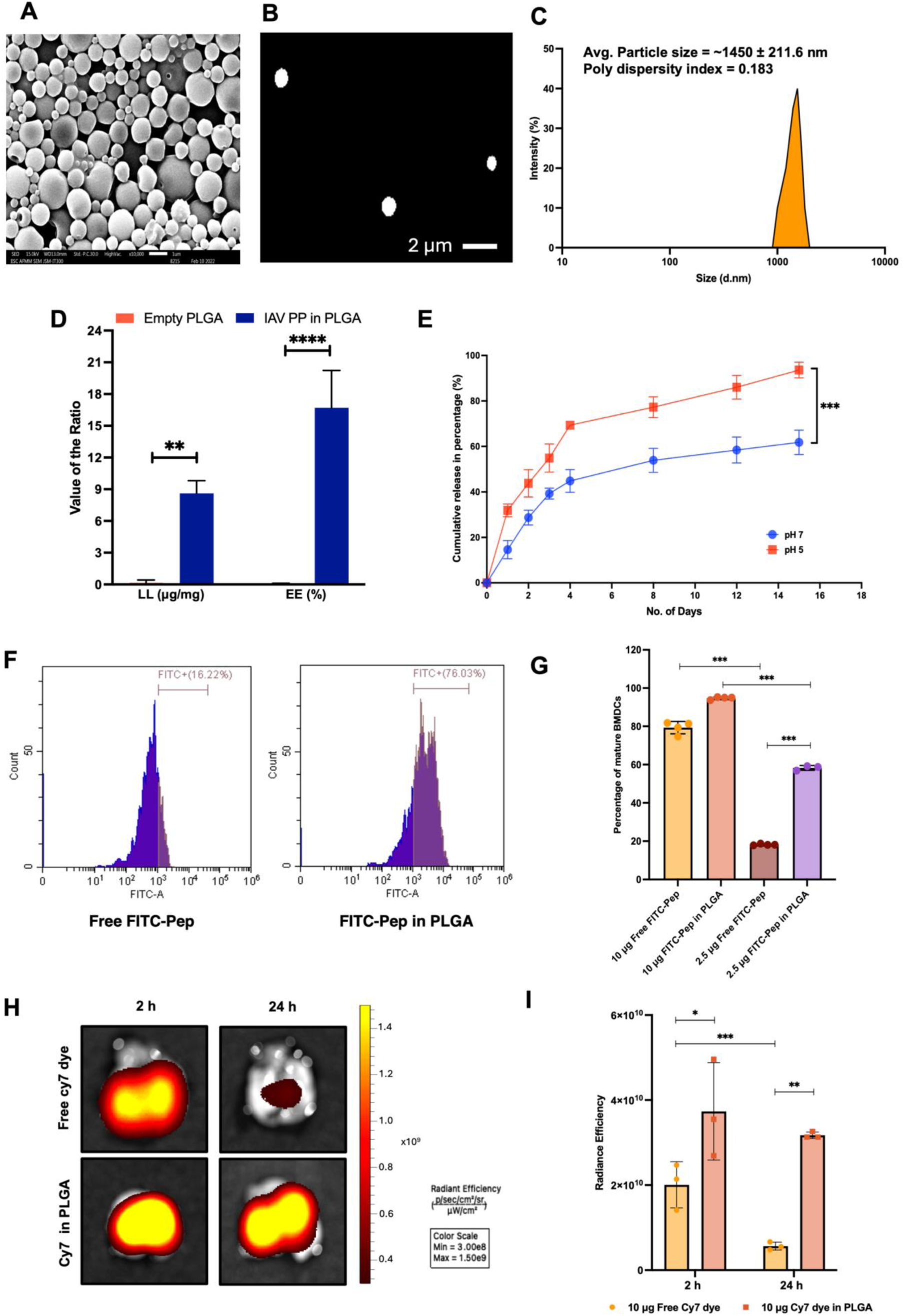
Structural and Functional Characterization of PLGA MPs loaded with CD8/CD4 bispecific IAV PP: **(A)** Size estimation through SEM, **(B)** Cy5-loaded PLGA-MPs imaging under 100x magnification in the fluorescence microscope, **(C)** size distribution of PLGA-MPs was determined using DLS. **(D)** Estimation of EE and LL using Fluorescamine assay **(E)** Analysis of pH-dependent release kinetics of PLGA-MPs, **(F, G)** FACS analysis of FITC-tagged PLGA-MPs uptake by mature BMDC cells (n=4), **(H, I)** IVIS imaging of Cy7-loaded PLGA-MPs retention in mice lungs (n=3) at 2 and 24 h was quantified by radiance efficiency. Statistical differences were determined using two-way ANOVA followed by Šídák’s test (D, E) and one-way ANOVA with Tukey’s test (G, I), with p-values defined as *p < 0.05, **p < 0.01, ***p < 0.001.

### Intranasal delivery of PLGA-MPs enhances pulmonary retention and promotes MHC-I cross-presentation of IAV PP

Next, we aimed to verify whether PLGA-MPs facilitate sustained and controlled release of encapsulated cargo in *in vivo* settings (*22, 26*). To ascertain this, we evaluated the retention of PLGA-MPs carrying a Cy7 fluorescent dye, serving as a surrogate for peptide cargo, in the lungs following intranasal delivery in mice. The *in vivo* imaging system (IVIS) used to track the intranasal delivery of Cy7-labeled PLGA-MPs to mouse lungs showed markedly different lung retention compared with the administration of the free Cy7 dye. The PLGA-encapsulated Cy7 exhibited prolonged lung retention, with strong fluorescence signals detectable for up to 24 h after administration (Figure 3H-I). In contrast, free Cy7 dye was rapidly cleared from the lungs, with minimal fluorescence detected by 24 h post-delivery. This demonstrates that PLGA encapsulation substantially enhances the residence time of cargo molecules in lung tissue, likely due to its larger size and the sustained release properties of the polymeric carrier. The improved retention of PLGA-MPs in the lungs suggests this delivery system could enhance the efficacy of intranasal vaccines by prolonging antigen exposure to immune cells in the respiratory tract.

Exogenously provided antigens, such as peptides in PLGA-MPs, are generally presented by MHC-II and activate a CD4+ T cell response (*26*). However, our peptide screening data indicated potent CD8+ T cell activation, in addition to CD4+ T cell activation, suggesting cross-presentation of processed antigens. To confirm this observation, we adapted the immunopeptidome method (*27*), wherein we incubated activated mouse BMDCs with PLGA-MPs carrying IAV PPs. The workflow involved immunoprecipitating MHC I molecules, followed by acid elution of bound peptides and mass spectrometry analysis (Supplementary Fig. 4A).

The resulting spectrum revealed multiple distinct peaks corresponding to known influenza-derived CD8+ T cell epitopes, which were covered by the CD8/CD4 bispecific peptides packaged in the PLGA-MPs (Supplementary Fig. 4B-C). These results demonstrate that PLGA-MP-based delivery effectively enables cross-presentation of exogenous IAV peptides, supporting its potential for eliciting cytotoxic CD8+ T cell responses and the expected helper CD4+ T cell response. To evaluate intracellular trafficking and endosomal escape, the localization of FITC-labelled peptide in THP1 cells was examined using confocal microscopy after treatment with either free FITC-peptide or PLGA-encapsulated peptide. As shown in Supplementary Fig. 3D, free FITC-peptide exhibited strong colocalization with lysosomal staining, indicating confinement within endolysosomal compartments. In contrast, PLGA-MP treated cells showed a more cytoplasmic FITC signal with reduced overlap with lysosomes, suggesting endosomal escape. Quantitative analysis confirmed these observations (Supplementary Fig. 4E-F), showing that the Manders’ correlation coefficient between FITC and lysosomes was significantly lower for the PLGA-MP-delivered peptide than for the free peptide.

### PLGA-MP encapsulation of IAV CD8/CD4 bispecific peptides enhances their T cell activation potential

Based on PLGA-MP characterization, the particles are expected to enhance APC uptake and enable sustained, pH-dependent intracellular antigen release, thereby improving antigen presentation and T cell activation. To evaluate this, we performed ex vivo T cell activation and proliferation assays using splenocytes from H1N1 (maCal/09 and X31) and H3N2 (HK/19) infected mice (Figure 4A). Cells were stimulated with 10 µg free IAV PP, 2.5 µg free IAV PP, 2.5 µg IAV PP encapsulated in PLGA-MPs, or 0.2 mg empty PLGA-MPs. T cell activation was assessed by measuring IFN-γ+ and TNF-α+ frequencies in CD4+ and CD8+ T cells (Supplementary Fig. 5A-B). Encapsulated IAV PP (2.5 µg) induced activation comparable to 10 µg free IAV PP in both H1N1 and H3N2 groups. Specifically, 2.5 µg IAV PP-loaded PLGA-MPs elicited CD4+IFN-γ+, CD4+TNF-α+, CD8+IFN-γ+, and CD8+TNF-α+ responses equivalent to 10 µg free peptide (Figure 4B, 4C, Supplementary Fig. 6A). Empty PLGA-MPs induced minimal activation, confirming antigen specificity. Enhanced activation was also observed in lung-resident CD4+ and CD8+ T cells from maCal/09- and X31-infected mice (Supplementary Fig. 6A, 6B, and Supplementary Fig. 7E, 7F). Given that the lung is the primary site of IAV infection and early replication, the presence of IFN-γ and TNF-α producing T cells indicates effective local cellular immunity. These results demonstrate that PLGA-encapsulated PP promotes both systemic and mucosal immune priming.

**Figure 4.**
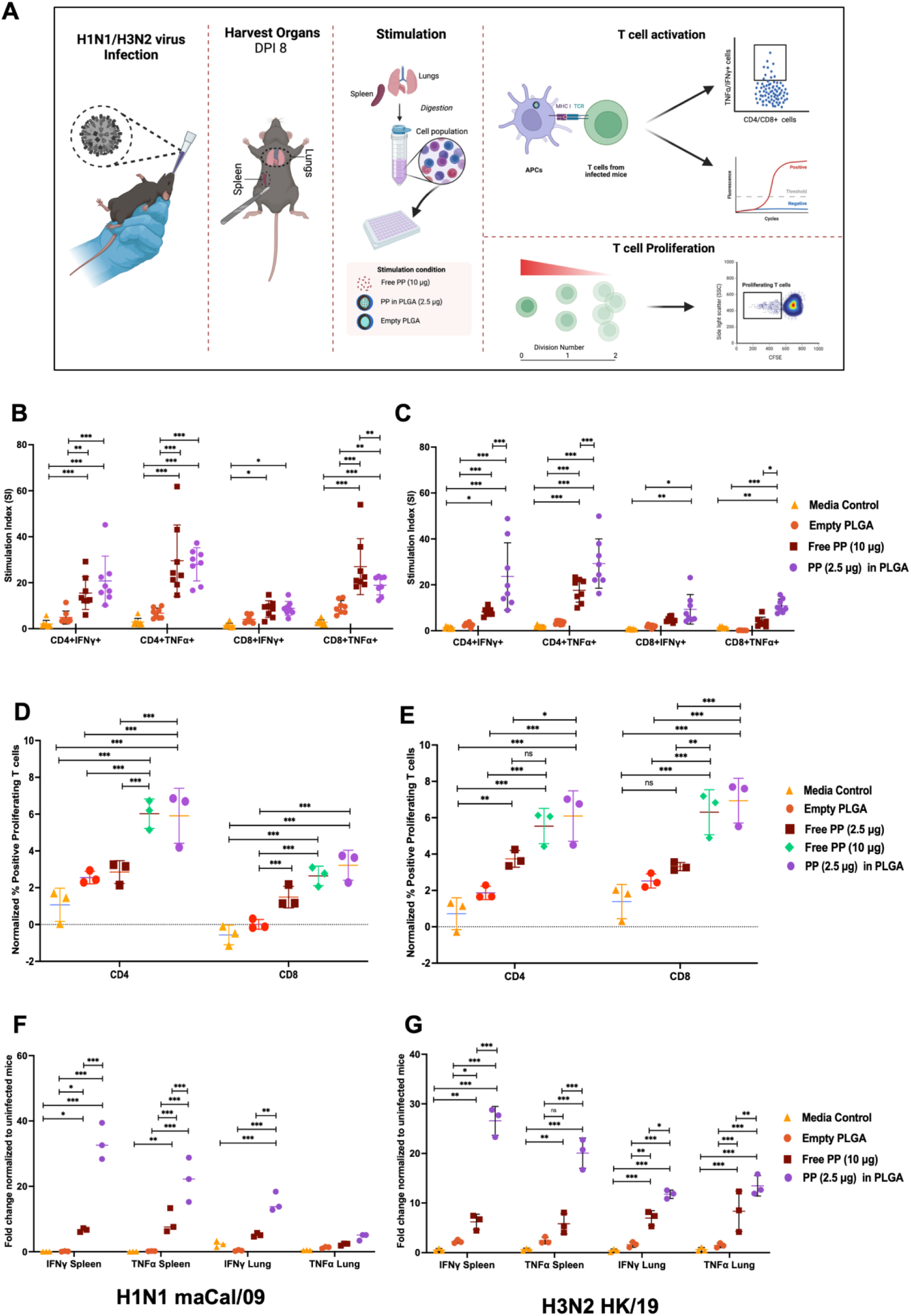
Comparing the T cell activation and proliferation of Free IAV PP and IAV PP Loaded in PLGA-MPs: The experimental workflow **(A)** evaluates T cell responses in ma/Cal/09 H1N1 **(B, D, F)** and HK/19 H3N2 **(C, E, G)** infected mice following splenocyte stimulation with various IAV PP formulations. Cytokine production (IFN-γ, TNF-α) was quantified by flow cytometry **(B, C)** and mRNA expression **(D, E)**, while proliferation was assessed by CFSE assay **(F, G)**. Data represent mean ± SEM, statistical significance was determined by one-way ANOVA with Tukey’s test. (*p < 0.05, **p < 0.01, ***p < 0.001).

**Figure 5.**
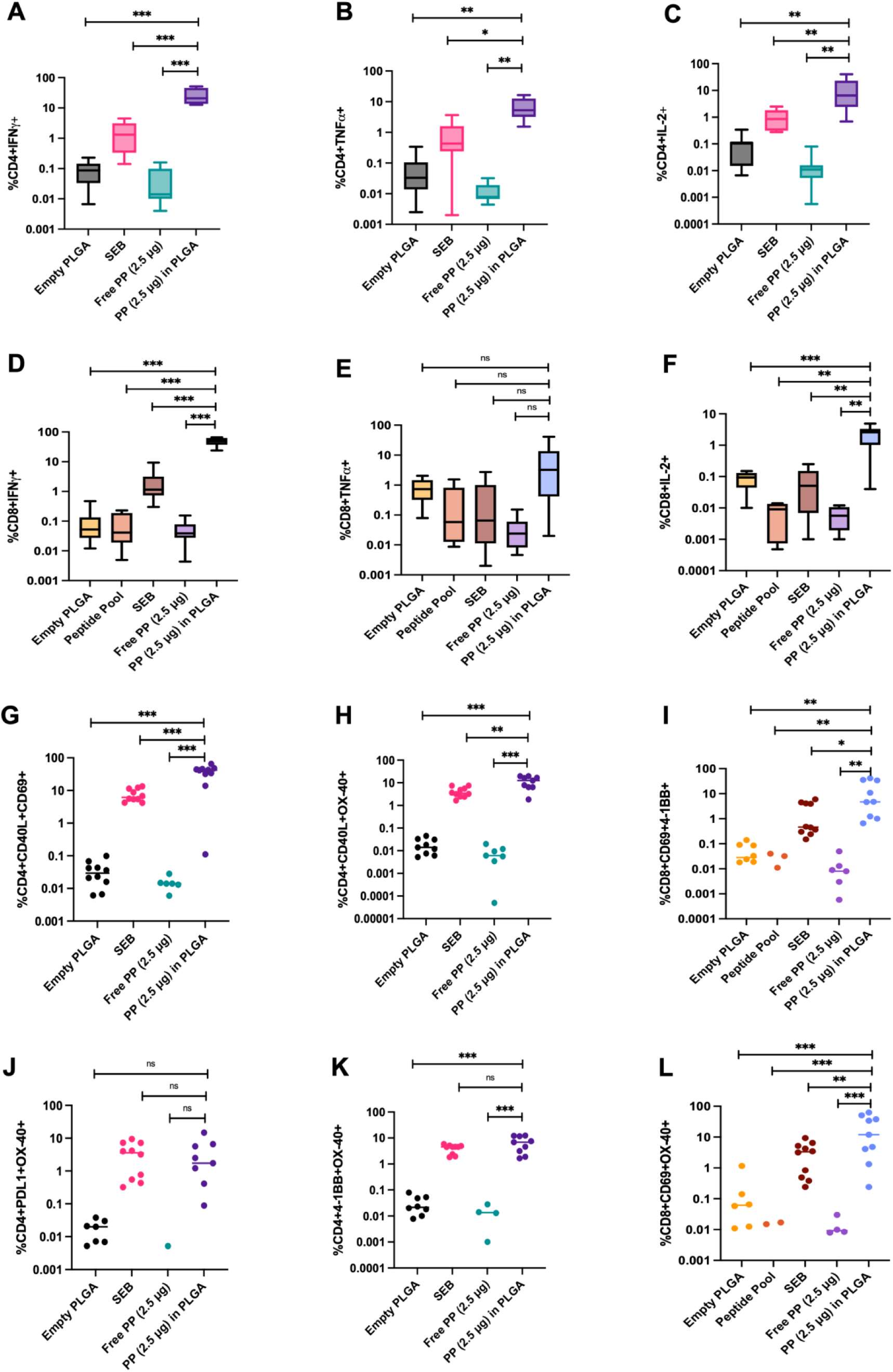
Activation and cytokine production of T cells in human PBMCs after stimulation with different vaccine formulations: (A-F) Percentage of cytokine-producing T cells. **(A-C)** show IFN-γ+, TNF-α+, and IL-2+ cytokine production, respectively, within the CD4+ T cell population and **(D-F)** within the CD8+ T cell population. **(G-L)** Quantification of activation-induced markers on T cells expressing canonical markers CD69, CD40L, OX-40, 4-1BB, and PD-1 was analyzed by flow cytometry. Each point represents an individual human sample; bars indicate mean ± SEM. Statistical significance was calculated using ordinary one-way ANOVA with Tukey’s multiple comparison test. (*p < 0.05, **p < 0.01, ***p < 0.001, ns: not significant).

**Figure 6.**
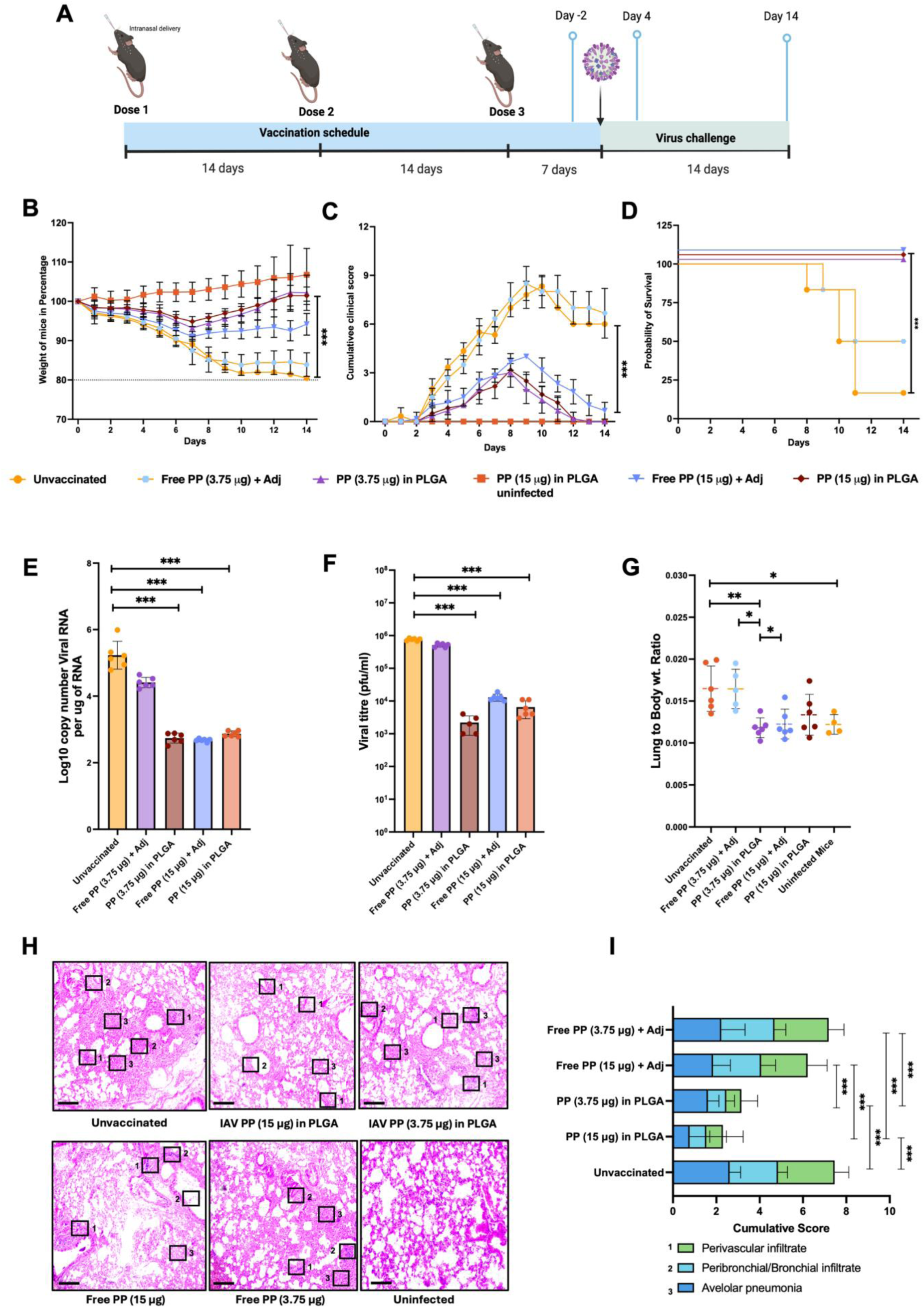
Evaluation of immune protection induced by intranasal vaccination against pandemic ma/Cal/09 H1N1 viruses: **(A)** Schematic for vaccination and immunization schedule followed for the study. **(B)** Post-virus challenge weight was monitored over 14 days. **(C)** Cumulative clinical scores, and **(D)** Survival rates. Vaccine efficacy was further evaluated by quantifying viral RNA copy number **(E)** and viral titer pfu/mL **(F)** in the nasal turbinate. **(G)** Lung-to-body weight ratios to assess pulmonary inflammation. **(H)** H and E staining images **(I)** Cumulative scoring measured perivascular infiltrate, bronchiolar infiltrate, and alveolar pneumonia severity. Data (n=6) are presented as mean ± SEM, with significance determined by a two-way ANOVA test. *p < 0.05, **p < 0.01, ***p < 0.001 and Scale bar = 200 µM

Importantly, PLGA-MP delivery induced polyfunctional CD4+ and CD8+ T cells co-expressing IFN-γ and TNF-α (Supplementary Fig. 5C, 5D and Supplementary Fig. 7B), highlighting enhanced functional quality at a reduced antigen dose. RT-PCR analysis further showed that splenocytes and lung cells stimulated with 2.5 µg encapsulated IAV PP expressed IFN-γ and TNF-α mRNA levels comparable to those induced by 10 µg free peptide in both H1N1 and H3N2 groups (Figure 4D, 4E, Supplementary Fig. 7C), whereas empty PLGA-MPs showed negligible cytokine expression. Furthermore, CFSE-based proliferation assays (Figure 4F, 4G, Supplementary Fig. 7D) demonstrated significantly increased CD4+ and CD8+ T cell proliferation following stimulation with encapsulated or free IAV PP compared to media and empty PLGA controls. Notably, 2.5 µg IAV PP-loaded PLGA-MPs induced proliferation comparable to 10 µg free peptide and significantly higher than 2.5 µg free peptide (Supplementary Fig. D). Overall, PLGA-MP encapsulation significantly enhances T cell activation, proliferation, and cytokine production, resulting in responses comparable to those with higher free antigen doses. These findings support PLGA-MPs as an efficient, dose-sparing antigen delivery platform with strong potential for vaccine development.

### PLGA-MP-encapsulated IAV-PP elicits robust T cell activation in PBMCs from vaccinated human subjects

So far, we have demonstrated that CD8/CD4 bispecific peptides derived from IAV consensus NP and M1 proteins activate mouse helper and cytotoxic T cells, and this activation is further enhanced when peptides are delivered using PLGA-MPs. To confirm whether human T cells also respond to IAV PP-loaded PLGA-MPs, we used peripheral blood mononuclear cells (PBMCs) from 5 subjects vaccinated with the trivalent IAV Vaccine (Table S4). This vaccine is enriched in IAV surface glycoproteins but also contains limited NP and M1 components, triggering the corresponding T cell response (*28*).

We stimulated them in vitro with CD8/CD4 bispecific peptide formulations (free PP vs. IAV PP loaded in PLGA-MP), and the resulting immune responses were characterized using flow cytometry to quantify cytokine-producing T cell subsets. Figure 5A-F illustrates the percentages of CD4+ and CD8+ T cells producing IFN-γ, TNF-α, and IL-2 cytokines. Among CD4+ T cells, the production of IFN-γ (A), TNF-α (B), and IL-2 (C) was significantly elevated in the group treated with 2.5 µg of IAV PP encapsulated in PLGA-MPs compared to both the unstimulated controls and the free PP (2.5 µg) condition. Similarly, CD8+ T cells demonstrated enhanced production of IFN-γ (D) and IL-2 (F) in response to the PLGA-formulated antigen. However, TNF-α (E) levels did not show a statistically significant increase. To further delineate T cell activation status, we assessed the expression of several activation-induced surface markers in Figure 5G-L. Specifically, we analyzed the expression of CD40L+, CD69+, OX-40+, 4-1BB+, and PD-L1+ on both CD4+ and CD8+ T cells. These markers offer insight into the functional activation and costimulatory potential of T cells following antigen stimulation. The increased expression of CD40L and OX-40 on CD4+ T cells suggests enhanced helper T cell activation and a potential for supporting B cell responses (Figure 5H). Upregulation of CD69, an early activation marker, was observed in both T cell subsets (Figure 5G, 5I, and 5L), indicating rapid immune activation.

CD69 appears rapidly upon activation and is involved in the retention of cells in lymphoid tissue. Additionally, elevated expression of 4-1BB on CD8+ T cells supports a cytotoxic memory response; it’s a costimulatory molecule on activated T cells (Figure 5I). PD-L1 on T cells indicates suppression of immune responses. Interestingly, PD-L1 expression patterns were also examined to assess potential regulatory feedback mechanisms induced by vaccination (Figure 5J). These results suggest that IAV PP loaded in PLGA-MPs (2.5 µg PP) induces robust activation of both CD4+ and CD8+ T cells, characterized by enhanced cytokine production and increased expression of activation markers. With validation of the mouse and human T cell activation potential of the CD8/CD4 bispecific peptides and loaded PLGA-MPs, we were ready to test the efficacy of this vaccine formulation in the murine model.

### Intranasal vaccination with IAV PP-loaded PLGA-MPs restricts pandemic H1N1 virus replication and pathology in a murine model

Following confirmation of robust T cell activation, we evaluated IAV PP-loaded PLGA-MPs as a vaccine formulation in the C57BL/6 mouse model. Intranasal delivery ensured deep lung deposition and sustained antigen release, supporting mucosal immunization. Given their APC-targeting size range and controlled release kinetics, PLGA-MPs were expected to possess intrinsic self-adjuvanting properties; thus, no external adjuvant was used for encapsulated peptides, whereas alum was included for free peptide controls. Two antigen doses (3.75 µg and 15 µg) were tested to assess dose dependency (Figure 6A). Mice received three intranasal immunizations at 14-day intervals. Two days before the challenge, spleen and lung tissues were analyzed for antigen-specific T cell responses using a BMDC-CD3+ T cell co-culture system with intracellular cytokine staining (Supplementary Fig. 8A-B). PLGA-encapsulated IAV PP induced significantly higher frequencies of IFN-γ+ and TNF-α+ CD4+ and CD8+ T cells compared to unvaccinated and free peptide groups. Notably, the 3.75 µg PLGA group generated robust cytokine responses in both spleen and lung, clearly outperforming the equivalent free peptide dose and approaching responses seen with 15 µg formulations. In the lungs, TNF-α+ T cells were particularly enriched in the 3.75 µg PLGA group, indicating strong mucosal activation. Proliferation assays further highlighted the advantage of encapsulation. In the spleen, both 15 µg and 3.75 µg PLGA groups showed marked CD4+ and CD8+ T cell expansion, whereas free peptide induced minimal proliferation (Supplementary Fig. 8C). In the lung, proliferation was predominantly observed in CD8+ T cells, with the 3.75 µg PLGA group demonstrating substantial expansion despite the lower antigen dose (Supplementary Fig. 8D). These findings indicate that even at one-quarter of the higher dose, PLGA encapsulation sustains strong systemic and mucosal T cell responses.

Protective efficacy was assessed following challenge with mouse-adapted maCal/09 H1N1 or X31/PR8 H1N1 viruses (Figure 6A, Supplementary Fig. 9A). Unvaccinated mice showed progressive weight loss beginning on Day 4, reaching ∼80% baseline by Day 14 (Figure 6B, Supplementary Fig. 9B). In contrast, the 3.75 µg PLGA group maintained significantly higher body weight throughout infection, comparable to or indistinguishable from the 15 µg groups. Clinical scores peaked sharply in unvaccinated animals but remained consistently low in the 3.75 µg PLGA group, demonstrating effective disease control (Figure 6C, Supplementary Fig. 9C). Survival analysis over 14 days revealed 70-80% mortality in unvaccinated mice and only 50% survival in the 3.75 µg free peptide group. Strikingly, the 3.75 µg PLGA formulation conferred 100% survival, matching the 15 µg groups and clearly outperforming the equivalent free peptide dose (Figure 6D, Supplementary Fig. 9D). This demonstrates superior protective efficacy at a substantially reduced antigen dose.

Assessment of viral replication showed that the 3.75 µg PLGA group significantly reduced viral RNA copies and infectious titers in the nasal turbinate, lungs, and nasal washes compared to the unvaccinated and 3.75 µg free peptide groups (Figure 6E, 6F, Supplementary Fig. 9E, F, Supplementary Fig. 10A, B). Viral suppression in the 3.75 µg PLGA group was comparable to that achieved with 15 µg formulations, indicating potent control of viral replication even at the lower dose. Inflammation analysis further supported its efficacy. The lung-to-body weight ratio at 4 DPI was significantly elevated in unvaccinated and 3.75 µg free peptide groups but markedly reduced in the 3.75 µg PLGA group (Figure 6G, Supplementary Fig. 9G). Histopathological examination revealed severe perivascular infiltration, peri-bronchial inflammation, and alveolar pneumonia in unvaccinated mice. In contrast, the 3.75 µg PLGA group showed substantially reduced cumulative pathology scores, with preserved lung architecture and minimal inflammatory damage (Figure 6H, 6I, Supplementary Fig. 9H, I). To evaluate dose sparing, mice received either two or three intranasal doses of 3.75 µg PLGA-encapsulated IAV PP before lethal challenge (Supplementary Fig. 11A). Both regimens prevented significant weight loss, enabled rapid recovery by Day 6, and achieved 100% survival (Supplementary Fig. 11B-C), with no observable difference between two and three doses.

Collectively, these results demonstrate that the 3.75 µg PLGA-MP formulation delivers superior performance: it induces strong mucosal and systemic T cell responses, markedly suppresses viral replication, minimizes lung inflammation and pathology, and provides complete protection against lethal influenza challenge. Importantly, this protection is achieved at a low antigen dose and with only two administrations, underscoring the potency and dose-sparing advantage of PLGA-based intranasal vaccination.

### Intranasal vaccination with PLGA-MPs restricts homologous H3N2 and heterologous H5N1 IAV replication and pathology in mice

To evaluate the breadth of protection conferred by PLGA-MP encapsulated IAV peptides, mice were immunized intranasally with 3.75 µg IAV PP in PLGA-MPs using the same three-dose regimen (Figure 7A), followed by challenge with seasonal A/Hong Kong/2671/2019 H3N2 (HK/19) and heterologous A/Vietnam/1203/2004 H5N1 (HALo) viruses. Clinical parameters, viral burden, lung inflammation, and survival were monitored for 14 days. Unvaccinated mice exhibited marked weight loss after both H3N2 and H5N1 infection. In contrast, mice vaccinated with 3.75 µg IAV PP in PLGA-MPs maintained stable body weight throughout the observation period, indicating effective disease control against both subtypes (Figure 7B). Clinical scores were significantly elevated in unvaccinated groups. In contrast, vaccinated animals displayed minimal symptoms following either H3 or H5 challenge (Figure 7C). Kaplan-Meier survival analysis demonstrated complete protection in vaccinated mice, with 100% survival against both H3N2 and the highly pathogenic H5N1 challenge. In contrast, unvaccinated controls showed substantial mortality, particularly following H5N1 infection (Figure 7D). Thus, the 3.75 µg PLGA-MP formulation conferred robust cross-subtype protection.

**Figure 7.**
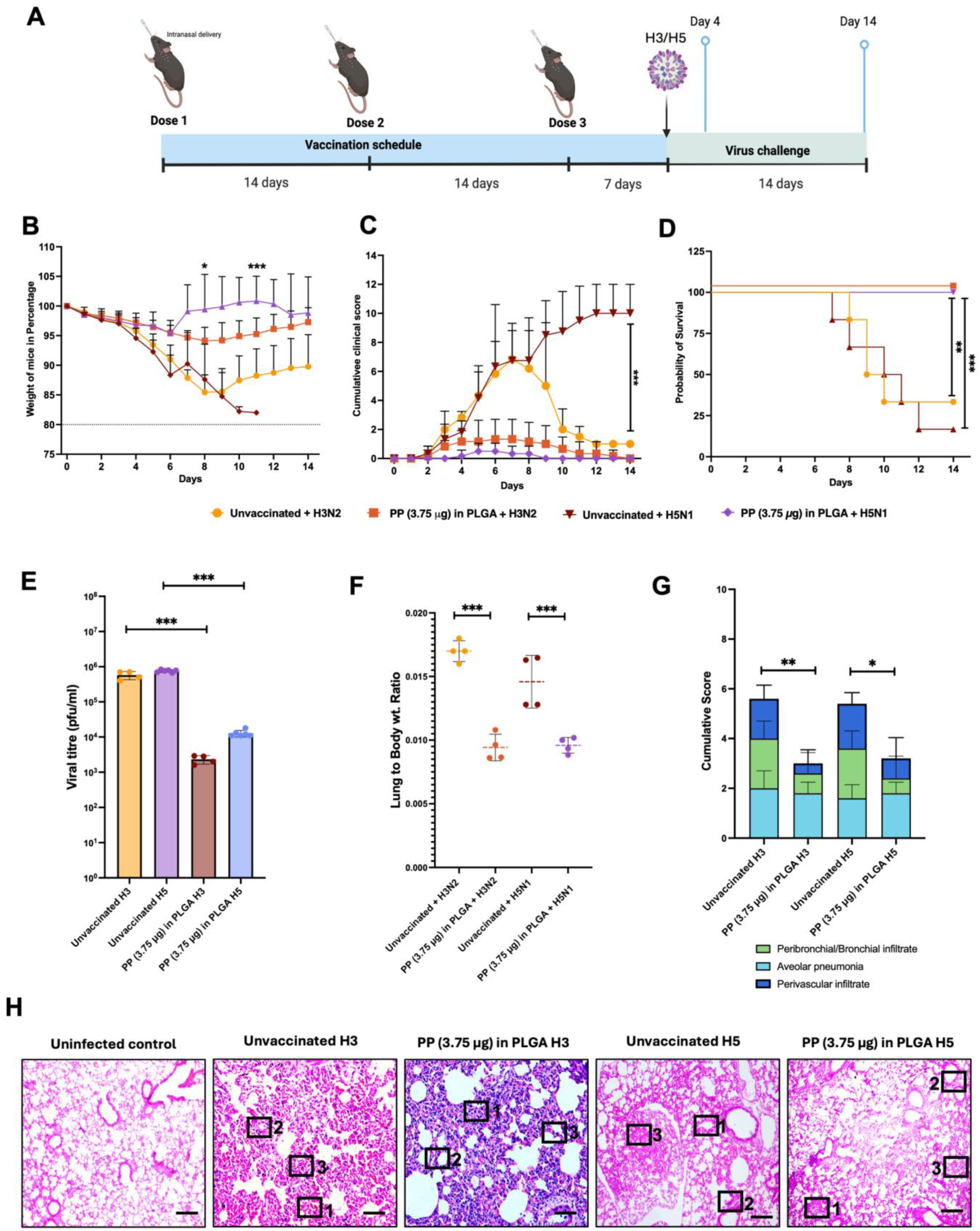
Assessment of heterosubtypic immunoprotection via intranasal vaccination against seasonal HK/19 H3N2 and avian HALo H5N1 viruses: **(A)** Schematic for the vaccination and challenge experiment conducted on mice. Post-virus challenge monitoring for 14 days included weight change **(B),** Cumulative clinical scores **(C),** and Survival graph **(D).** Vaccine efficacy was evaluated by quantifying viral titers in the nasal turbinate via plaque assay **(E)** and **by** lung-to-body weight ratio **(F)** to assess pulmonary inflammation. Histopathological analysis of H&E-stained lung tissues **(H)** provided cumulative scoring for perivascular, bronchial, and alveolar pneumonia severity **(G)**. Data (n=4-6) are presented as mean ± SEM. Statistical significance was determined by a two-way ANOVA test. *p < 0.05, **p < 0.01, ***p < 0.001 and Scale = 200 μm.

Viral titers measured by plaque assay in nasal turbinate tissue, lungs, and nasal washes showed significantly reduced viral replication in vaccinated mice compared to unvaccinated controls (Figure 7E, Supplementary Fig. 12A, B). Unvaccinated animals exhibited the highest lung viral loads, whereas the 3.75 µg PLGA group achieved pronounced suppression of infectious virus across upper and lower respiratory tissues, demonstrating strong antiviral efficacy. Pulmonary inflammation assessed by lung-to-body weight ratio at 4 DPI was significantly elevated in unvaccinated mice but markedly reduced in vaccinated animals (Figure 7F). Histopathological analysis further confirmed severe bronchial infiltration, alveolar damage, and hemorrhagic lesions in unvaccinated mice. In contrast, vaccinated groups showed minimal inflammatory infiltration and preserved alveolar architecture (Figure 7G-H). Cumulative pathology scores were substantially lower in vaccinated mice, indicating protection from virus-induced lung injury. Collectively, these findings demonstrate that intranasal vaccination with 3.75 µg PLGA-MP-encapsulated CD8/CD4 bispecific peptides derived from conserved NP and M1 proteins induces broad, cross-subtype protection against both H3N2 and H5N1 viruses. The formulation effectively prevents weight loss, reduces viral replication, limits lung inflammation and tissue damage, and ensures complete survival, highlighting its capacity to confer robust and broadly protective T cell-mediated immunity.

## Discussion

The development of broadly protective influenza vaccines remains a central challenge in infectious disease research, driven by the rapid antigenic evolution of IAVs (*34*). Antigenic drift, driven by the error-prone RNA-dependent RNA polymerase (RdRp), enables viral evasion of pre-existing immunity (*30*). Genetic reassortment between IAV strains can cause antigenic shift, occasionally leading to pandemic viruses (*31*). Current vaccine strategies predominantly target variable surface glycoproteins, such as HA and NA, which are subject to immune selection pressure and often fail to provide optimal protection due to antigenic mismatch. At the same time, substantial efforts for broadly protective vaccines focus on conserved humoral targets, including the HA stalk, chimeric or consensus HA constructs, conserved regions of neuraminidase (NA), and the M2 ectodomain (*32, 33*). The HA stalk is relatively conserved but has limited ability to elicit neutralizing antibodies (*45*). Computational approaches, such as COBRA-designed consensus HA, have shown promise (*14*). Alternatively, vaccines targeting conserved internal proteins NP and M1 aim to induce cross-reactive immunity (*35*).

Although humoral immunity is widely regarded as the primary correlate of protection against IAV infection, accumulating evidence highlights the role of cellular immunity in viral clearance and disease severity (*36, 37*). Human challenge studies demonstrate that influenza-specific T cell responses can mitigate clinical illness, as individuals lacking pre-existing neutralizing antibodies but possessing T cell immunity show reduced viral shedding following infection (*38*). Studies revealed that both CD4+ and CD8+ T cells contribute to viral clearance and reduced disease severity (*39*). Challenge studies using pandemic H1N1 virus established a strong inverse correlation between pre-existing CD8+ T cell immunity and viral load, symptom severity, and illness duration, along with marked expansion of virus-specific CD8+ T cells after infection (*40*). Historically, CD8+ T cells were considered the dominant mediators of cellular immunity against IAV, supported by studies in mice lacking functional CD8+ T cells (*10*). However, emerging evidence highlights a critical and previously underappreciated role for CD4+ T cells in protection against lethal influenza infection (*41, 42*). Supporting this, murine studies show that depletion of either CD4+ or CD8+ T cells alone does not abolish protection, whereas simultaneous depletion of both subsets prevents viral clearance (*43*). Moreover, in non-lethal IAV infections, CD8+ T cells appear dispensable for viral clearance (*42*), emphasizing the importance of additional immune mechanisms, particularly CD4+ T cell-mediated responses, in controlling infection.

A key limitation of existing T cell-based vaccine approaches is their reliance on individual epitopes, which often exhibit limited stability, restricted HLA coverage, and suboptimal induction of coordinated immune responses. Here, we address this challenge by targeting viral protein regions enriched with multiple overlapping, immunodominant CD4+ and CD8+ T cell epitopes, rather than focusing on individual CD8+ T cell epitopes, thereby eliciting a broader yet stronger T cell response against IAV infection. To test this idea, we mapped CD8/CD4 bispecific regions within the NP and M1 consensus sequences of H1N1 and H3N2 IAV strains. These consensus sequences were generated using a computational micro-consensus approach similar to COBRA (*14*). However, unlike the conventional COBRA method, we constructed the consensus using protein sequences instead of nucleotide sequences and incorporated all available complete human isolate sequences reported since 1918. This strategy produced a micro-consensus with improved homology across diverse IAV evolutionary lineages compared with traditional consensus sequences. NP and M1 were selected as vaccine targets because they are produced in large quantities during infection, increasing the likelihood that their peptides will be processed, presented, and recognized by T cells. A major limitation of bioinformatic methods for identifying CD8+ and CD4+ T cell epitopes is the generation of false-positive predictions. To overcome this problem, we used a combinatorial prediction strategy using three widely used tools (IEDB, ProPred, NetMHCpan) (*16–18*). Epitope selection prioritized broad human HLA coverage and conservation across IAV strains. Notably, all 16 selected NP and M1 regions contained both human-specific and mouse-specific CD4+ and CD8+ epitopes, enabling evaluation in mice. Furthermore, the CD8/CD4 bispecific activity of the six best-performing peptides was validated using human PBMCs from vaccinated subjects. Notably, these peptides induced stronger T cell activation in human cells than in mouse cells, likely because the epitope prediction pipeline was optimized using human epitope databases.

T cell peptide-based vaccines face several limitations, including poor peptide stability, a short in vivo half-life due to rapid clearance from the respiratory mucosa via mucociliary activity and systemic absorption, inefficient delivery to APCs, and a requirement for adjuvants (*44*). To address these challenges, we employed PLGA-MPs as a delivery platform. PLGA is a biodegradable polymer widely used in FDA-approved clinical applications (*45*). Encapsulation of peptides within PLGA-MPs protects them from extracellular proteolytic degradation (*26*). The PLGA-MPs used in this study ranged in size from 1-5 µm, a size range known to facilitate efficient uptake by APCs (*24*). From a formulation perspective, optimizing peptide encapsulation efficiency is critical. One strategy to improve encapsulation involves ion pairing of peptides with oppositely charged counter-ions before particle fabrication (*46*). This approach increases peptide hydrophobicity, limits premature diffusion into the aqueous phase during preparation, and improves peptide loading and retention within the polymer matrix. Such strategies may increase encapsulation efficiency to above 70-80%, reducing peptide loss and enabling scalable production (*46*). In our study, PLGA MP-encapsulated peptides showed higher uptake by mouse BMDCs than free peptides. The MPs also displayed sustained, pH-dependent release, supporting prolonged immune engagement and intracellular peptide release within APCs.

Protein subunit and peptide vaccines generally require adjuvants to generate a depot effect and stimulate innate immune responses (*47*). In contrast, the particulate nature of PLGA-MPs and their capacity to disrupt plasma and endosomal membranes can directly activate innate immune pathways through pattern-recognition receptors, including Toll-like receptors (*1*). This activation promotes recruitment and maturation of APCs, thereby enhancing T cell priming. Leveraging these intrinsic immunostimulatory properties, we developed a PLGA-MP based vaccine formulation without conventional adjuvants, thereby minimizing associated safety concerns. In preclinical studies, PLGA-MP encapsulated peptides showed superior efficacy compared with free IAV PP formulated with the traditional adjuvant Alhydrogel.

Current seasonal influenza vaccines are typically administered intramuscularly, primarily inducing systemic immunity (*50*). In contrast, our PLGA-MPs were safely delivered intranasally, eliciting both mucosal immunity in the upper and lower respiratory tract and systemic immune responses. This was reflected by T cell activation in lung tissue and splenocytes of vaccinated mice, along with a marked reduction in viral load in the nasal turbinate and lungs. Although exogenous peptides delivered via PLGA-MPs are typically expected to be presented via MHC-II to helper T cells, we observed a strong MHC-I dependent CD8+ T cell response, indicating efficient cross-presentation. PLGA-MPs have been reported to escape the endosomes by selectively changing the surface charge of MPs (from anionic to cationic) in the acidic endo-lysosomal compartment, which causes the MPs to interact with the endo-lysosomal membrane and deliver the cargo into the cytosol (*51*). This should also facilitate MHC-I cross-presentation of CD8 epitopes derived from the CD8/CD4 bispecific peptides loaded in the PLGA-MPs (*52, 53*). This was confirmed by an immunopeptidome assay (*27*), which revealed that BMDCs presented epitopes from all six CD4+/CD8+ bispecific peptides.

Compared with alternative platforms, including viral vectors such as adenovirus or modified vaccinia Ankara (MVA), mRNA-based vaccine platforms, this approach offers several advantages. Viral vectors can induce strong T cell immunity, but are often limited by strong anti-vector immune responses (*54*). Meanwhile, mRNA-based approaches are being actively explored for broadly protective influenza vaccines (*55*). Incorporating CD4+/CD8+ bispecific peptide components into mRNA constructs may further enhance their breadth and efficacy by strengthening cellular immunity. The concept of integrating humoral and cellular immunogens in a synthetic vaccine has previously been explored by BiondVax’s multimeric peptide vaccine, which contains conserved B- and T cell epitopes from both influenza A and B viruses and is formulated with the oil-based adjuvant Montanide ISA 51VG (*56*). Although this candidate has progressed to Phase 3 clinical trials, Montanide has been associated with safety concerns, including local injection site reactions and systemic inflammatory responses, which may limit its suitability for repeated dosing or use in vulnerable populations.

In contrast, our formulation avoids reactogenic oil-based adjuvants. Instead, it uses PLGA, a well-characterized, FDA-approved biodegradable polymer with an established safety profile, enabling intranasal delivery and the induction of mucosal immunity. PLGA degrades into lactic and glycolic acids, which are naturally metabolized, further supporting its safety and biocompatibility (*22, 45*). The development of an inhalable, lyophilized form of the PLGA-MP vaccine formulation to improve ease of use could further enhance translatability. Notably, we previously demonstrated that PLGA particles can be engineered into inhalable dry powder formulations that achieve efficient pulmonary deposition in murine models (*57*). Such a needle-free approach may also improve vaccine acceptance and uptake by enabling easier administration and the potential for self-delivery (*58*). It will be important to investigate whether our formulation can serve as a booster or be combined with conventional HA-based vaccines to enhance vaccine efficacy and broaden protection.

In conclusion, we demonstrate a novel T cell-focused vaccine strategy that provides broad protection against IAVs. This platform offers a versatile template that could be adapted to target other rapidly evolving pathogens. The promising preclinical results support further optimization and clinical evaluation, with the potential to contribute to the development of next-generation influenza vaccines that may provide broader, more durable protection against this major global health threat.

### Limitations of the study

This study provides preclinical proof-of-concept for a broadly protective T cell-based IAV vaccine, but several limitations remain. The cellular and molecular mechanisms underlying vaccine-mediated protection require further investigation, particularly in clinically relevant models. The breadth of protection must be validated against a larger, more diverse panel of IAV strains, and the durability of vaccine-elicited immune memory remains to be established. A more comprehensive safety assessment, including mucosal and systemic toxicity following intranasal delivery, will be required. Comparing the immunogenicity and efficacy of this vaccine with those of standard IAV vaccines would be useful. However, there is no clinically approved T cell-based vaccine to serve as a benchmark. Finally, the potential applicability of this platform to other rapidly evolving respiratory pathogens warrants further evaluation.

## MATERIALS AND METHODS

### Resource availability

#### Lead contact

Further information and requests for resources and reagents should be directed to and will be fulfilled by the lead contact, Shashank Tripathi.

#### Materials availability

This study did not generate new reagents.

### Experimental model and subject details

#### Ethics Statement

IAV challenges in mice were conducted at the Viral BSL-3 facility with ethics approval (IAEC/IISc/ST/748/2020) following the recommendations of the Indian Council of Medical Research (ICMR) and Department of Biotechnology (DBT). The experiments were conducted in accordance with the CPCSEA (Committee for Control and Supervision of Experiments on Animals) guidelines.

#### Animal experiments

Female C57BL/6 mice (6-8 weeks) from Central Animal Facility, Indian Institute of Science, Bengaluru, India were housed at in groups of five in individually ventilated cages, maintained at a temperature of 23 ± 1 °C and a relative humidity of 50 ± 5%. The animals were provided with standard pellet feed and water ad libitum and maintained on a 12-hour day/night light cycle at the Viral Biosafety Level-3 facility of the Indian Institute of Science. All animals were monitored daily during the experiment. An overdose of Ketamine (Bharat Parenterals Limited) and Xylazine (Indian Immunologicals Ltd) was used to sacrifice animals upon completion of the experiment.

#### Cell lines and viruses

Madin-Darby Canine Kidney (MDCK) (NCCS, Pune, India) cells were cultured in Dulbecco’s modified Eagle medium (12100-038, Gibco) while human monocytic THP-1 cells (ATCC, TIB-202) were maintained in Roswell Park Memorial Institute (RPMI) 1640 medium (11875-093, Gibco). Supplemented with 10% HI-FBS (16140-071, Gibco), Penicillin-Streptomycin (15140122, Gibco), and GlutaMAX (35050-061, Gibco). Cells were cultured at 37°C in a humidified atmosphere with 5% CO₂. Influenza A virus strains maCal/09 H1N1, HK/19 H3N2, X31 H3N2, and HALo H5N1 were propagated in embryonated chicken eggs and titrated by plaque assay in MDCK cells (*59*).

## Method details

### Protein Sequence Retrieval

Unique NP and M1 protein sequences of human-infecting H1N1 and H3N2 IAVs from 1918 to 2019 were retrieved from the Bacterial and Viral Bioinformatics Resource Centre (BVBRC) database (https://www.bv-brc.org/), formerly the fluDB database (https://www.fludb.org) (*60*). Complete sequences were selected, excluding those of incomplete, putative, and laboratory-cultured strains. The sequences were grouped into clusters based on their H1/H3 clade classification, as shown in Table 1.

### Prediction of T cell Epitopes

T cell epitopes for the NP and M1 proteins were predicted using the IEDB I/II (*16*), NetMHCpan I/II (*17*), and Propred I/II (*18*). These tools provided potential epitopes based on binding affinity. Epitopes were selected using a percentile rank cutoff (top 2% for MHC I and 10% for MHC II). We selected 112 experimentally validated T cell epitopes for further analysis.

### Chemical synthesis of selected bispecific T cell peptides

To perform functional evaluation of the selected bispecific peptides, we chemically synthesized 16 CD4+/CD8+ bispecific peptides from GenScript Biotech (Nanjing, Jiangsu, China), along with well-studied control CD8+ T cell epitope NP-SRYWAIRTR (*61*), M1-GILGFVFTL (*62*) and CD4+ epitope NP-QVYSLIRPNENPAHK (*63*), M1-SGPLKAEIAQRLEDV (*64*) epitopes that are present in both H1N1 and H3N2 viruses. Also, the FITC tagged peptide (AYERMCNILKGKFQTAAQRAMVDQ, P6). The peptides were >95% pure, as verified by high-performance liquid chromatography (HPLC) and mass spectrometry. Upon receipt, lyophilized peptides were reconstituted in dimethyl sulfoxide (DMSO) or PBS to prepare stock solutions, aliquoted, and stored at –80 °C.

### Preparation of PLGA-MPs entrapped conserved IAV peptides

PLGA-MPs (Mw 10-15 kDa, 50:50, Akina AP041) were synthesized via solvent evaporation. The six conserved H1/H3 peptides of NP and M1 protein (6 mg total) were suspended in PBS, mixed with 2% PVA, 2% sucrose, and 2% Mg(OH)2, and emulsified in 150 mg PLGA dissolved in 10 mL DCM by homogenization (12,000 rpm, 2 mins). For specific experiments, 25 μg/mL of Cy5 or Cy7 was added to the PLGA-DCM solution. The emulsion was combined with 2% PVA, homogenized, and stirred for 3-4 h for DCM evaporation. After centrifugation at 11,000×g, the pellet was washed, frozen at −80°C, and lyophilized at −45°C. The particles were suspended in PBS, UV-sterilized, and analyzed using dynamic laser light scattering.

### Scanning Electron Microscopy

Scanning electron microscopy (SEM) was used to examine particle morphology. Particles were suspended in distilled water (0.5 mg/mL) and sonicated. 10 μL of the sample was deposited on carbon tape affixed to a metal stub and vacuum-dried. The sample was sputter-coated with gold (20 nm) using Bal-tec SCD 500, then analyzed using ThermoFisher XL-30 ESEM microscope with an electron beam (5 keV).

### Estimating the encapsulation efficiency

A standard curve was prepared using Bovine serum albumin (0-500 µg/mL) in PBS to estimate IAV peptides in PLGA particles. IAV peptide encapsulation was determined by dissolving 10 mg of PLGA particles in 100 μL DMSO, diluting 10-fold with distilled water, and centrifuging at 11,000 g for 10 minutes. The supernatant was collected, and Fluorescamine dye (1 mM in acetone) was added; the mixture was incubated for 5 minutes, and fluorescence was measured at 400/460nm using a Tecan Microplate Spectrophotometer. Loading efficiency (LE) refers to the actual amount of IAV PP incorporated per unit dry weight of PLGA-MPs. Encapsulation efficiency (EE) represents the percentage of the total peptide used during formulation that is successfully entrapped within the PLGA-MPs and is calculated as (amount of IAV PP encapsulated / total amount of IAV PP added) × 100%.

### pH-dependent release profile of PLGA-MPs

IAV PP-loaded PLGA-MPs (1 mg/mL) were incubated in PBS (pH 7) and MES buffer (pH 5) at 37 °C. At various time points, particles were pelleted at 11,000 × g and dissolved in 100 μL DMSO. The released peptide was quantified using the CBQCA Plus protein kit (Invitrogen, A66522) per the manufacturer’s instructions. Tecan Microplate Spectrophotometer measured fluorescence (465/550 nm). IAV peptide release was determined using the IAV PP standard curve in triplicate.

### Murine Bone Marrow-Derived Dendritic cells (BMDCs) isolation and culture

C57BL/6 WT mice (8-10 weeks) were euthanized, and the femur and tibia were dissected and treated with 70% ethanol for 5 minutes. The bone ends were cut, and the media was passed through a 26-G needle into a sterile dish. Cells were passed through a 19-G needle to break clumps. After RBC lysis, cells were seeded at 2 × 10^6^ cells/well with 20 ng/mL GM-CSF. Half the media volume was replaced with fresh GM-CSF media every other day (*65*). On day 7, cells were washed and incubated. BMDCs were characterized by CD11c+ (BD Biosciences, 564079), F4/80+ (BD Biosciences, 565411), CD80+ (17080182), CD86+ (Invitrogen, 47086282), and MHCII+ (Invitrogen, 1453282) markers using flow cytometry.

### Cellular Uptake of PLGA-MPs by mature BMDCs

To evaluate cellular uptake efficiency, mature BMDCs were seeded into 24-well plates. The cells were then incubated with either free FITC-labeled peptide or FITC-labeled peptide encapsulated in PLGA microparticles (PLGA-MPs). Two different peptide-equivalent doses were tested for each formulation: 2.5 µg and 10 µg. After a predetermined incubation period, the cells were washed thoroughly to remove any non-internalized peptides or microparticles. The percentage of cells that had taken up the peptide was quantified by measuring the FITC signal using flow cytometry. The results were expressed as the percentage of FITC-positive cells within the gated BMDC population.

### Confocal Imaging

THP-1 cells were seeded on coverslips and incubated with either free FITC-labelled peptide or PLGA-MP–encapsulated FITC-peptide at 10µg/ml concentrations for 3 h at 37°C. Cells were washed with PBS, fixed with 4% paraformaldehyde, permeabilized using 0.1% Triton X-100, and blocked with 3% BSA. Lysosomes were stained using anti-LAMP-1 antibody followed by a fluorophore-conjugated secondary antibody, and nuclei were counterstained with DAPI. Images were acquired using a confocal microscope under identical acquisition settings across conditions. Colocalization analysis was performed using ImageJ (Coloc2 plugin). Manders’ coefficients were calculated, where M1 represents the fraction of FITC (green) overlapping with LAMP-1 (red), and M2 represents the fraction of LAMP-1 overlapping with FITC. Cellular uptake was quantified as mean fluorescence intensity (MFI) of the FITC channel after background subtraction.

### BMDC-T Cell co-culture assay

From naïve mice, BMDCs were generated as described before (*66*). After 7 days, mature BMDCs were pulsed overnight with selected T-cell epitopes. Isolate spleen from vaccinated mice, enrich CD3 population using Mojosort^TM^ kit (Biolegend, 480023), and seed mature BMDCs at a 1:10 ratio and incubate for 48 hrs.

### Immunoprecipitation of MHC I molecule and mass spectrometry

Mature BMDCs from naïve mice were pulsed with 10μg IAV PP in PLGA-MPs for 6-8 hrs. After loading, BMDCs were lysed with mild buffer (1% NP-40, 150 mM NaCl, 20 mM Tris-HCl, pH 7.4, protease inhibitor). Lysates were clarified by centrifugation (14000xg, 15 mins, 4°C). MHC I complexes were immunoprecipitated using anti-MHC I antibody (clone IVA26, MA5-44043) with Protein G magnetic beads overnight at 4°C. Beads were washed, and peptides were acid-eluted using 10% acetic acid. Eluted peptides were subjected to liquid chromatography–mass spectrometry (LCMS) analysis. The acquired RAW data files were converted to mzML format and analyzed using MSGF+ (version 2024.03.26) (*67*) for peptide identification against the 6 flu antigen FASTA database. Searches were performed with unspecific enzyme settings to account for the non-tryptic nature of MHC-bound peptides, and appropriate precursor mass tolerance of 20 ppm and peptide length constraints (7–15 amino acids) were applied. The resulting peptide-spectrum matches were further processed and visualized using PeptideShaker (version 3.0.11) (*68*), enabling confidence scoring, filtering, and validation of identified peptides. Identified peptides were then mapped to their respective antigen sequences to determine MHC class I–associated epitopes.

### Immunization and Influenza Virus infection in a mouse model

Mice were immunized intranasally with 50 μL of 15 and 3.75 μg IAV PP with Alhydrogel (1:1 v/v) and IAV PP in PLGA. Two boosters were given at 2-week intervals. After final immunization, mice were anesthetized with Ketamine (60 mg/kg) and Xylazine (16 mg/kg) and challenged with viral suspensions: A/California/04/2009 H1N1 (ma/Cal/09, ∼50 pfu/ml), A/Hong Kong/2671/2019 H3N2 (HK/19, 1.5 x10^4^ pfu/ml), A/X-31 H3N2 (PR8, 200 pfu/ml) and A/Vietnam/1203/2004 H5N1 (HALo, 300 pfu/ml). Nasal turbinate tissue, lung tissue, and nasal washes were collected to determine viral titers by plaque assay and qRT-PCR.

### Residence time of PLGA MPs in Mouse lungs

Mice received intranasal administration of 50 µL containing 15 µg Cy7-labeled PLGA microparticles or 15 µg free Cy7 dye. IVIS imaging was performed immediately after administration and at 24 hrs (*69*). Explant mouse lungs were placed in the IVIS chamber, and fluorescence images were acquired using Cy7 filters. Living Image^TM^ software 4.8.2 (Revvity) was used to quantify radiance efficiency in the lungs, with average values calculated for each group at each time point.

### Isolation of splenocytes and lung cells from mice

Mice were euthanized by cervical dislocation on day 8 post-infection with the IAV virus. The spleen/lungs were extracted after sterilizing the abdominal area with 70% alcohol. The organs were rinsed with PBS and RPMI 1640, then placed in fresh RPMI 1640. Lungs were washed with PBS to remove blood. Spleen and lung cells were extracted using frosted glass slides. Lung tissue was incubated at 37°C for 30 mins in PBS containing 0.5% BSA, 5 mg/mL collagenase type 1, and 1 mg/mL DNase I. Samples were centrifuged at 800xg for 5 mins at 4°C, treated with RBC lysis buffer for 5 mins, washed with RPMI containing 0.5% BSA, filtered through a 70 μm Cell Strainer, and stored at −80°C in FBS containing 10% DMSO for FACS analysis.

### Intracellular Cytokine staining

Splenocytes from mice at 8 DPI or 4 DPV were stimulated with IAV PP for NP and M1 protein (10 μg/mL) for 9-12 h, with Brefeldin A (Ebioscience, 00450651) added. Cells were stained with viability dye (Thermo Fisher Scientific, L34966A) and antibodies: anti-CD3-APC-eFluor 780 (Invitrogen, 47038182), anti-CD4-FITC (Invitrogen, 11-0042-82), and anti-CD8-Super Bright 780 (Invitrogen, 78-0081-82) at 4°C for 30 mins. After washing and fixing with 4% buffered formalin (4°C, 20 mins), cells were permeabilized with 0.25% Saponin (Sigma Aldrich, 84510), then stained with anti-TNF-PE-Cy7 (Invitrogen, 25-7321-82), anti-IFN-γ-PE (Invitrogen, 12-7311-82), and anti-IL-2-APC (Invitrogen, 177021-82) at 4°C for 50 mins. Cells were analyzed on a Beckman Coulter Cytoflex Flow cytometer. Cumulative fold change was calculated by summing fold changes across cytokine-positive subsets, with frequencies background-corrected relative to unstimulated controls. For human PBMC T cell activation, 1-2x10^6^ cells were stimulated with IAV PP in PLGA-MPs (2.5 µg), Free IAV PP (2.5 µg), CD8 PP (2.5 µg) containing M1GILGFVFTL, NS1(122-130), PB1(413-421), NA(75-84), PA(225-233), PA(548-557), PA(46-54) HLA-A*02:01/03 peptides, and SEB (0.5 µg) for 2 h, followed by brefeldin A addition. Cells were stained with LIVE/DEAD dye and antibodies CD8-BV570, CD3 BUV805, and CD4-BV480, then permeabilized and stained with IFN-γ-PE-Cy7, TNFα-BB700, and IL-2-BV650. PLGA and unstimulated controls were used to set cytokine expression gates.

### Activation Induced Marker (AIM) assay

HLA-typed PBMCs were thawed and rested for 2 h at 37°C, 5% CO2, in 10% complete RPMI. Cells (1x10^6^) were plated, and the CD40 blocking antibody was added for 15 minutes at 0.5 μg/mL. PBMCs were stimulated with IAV PP in PLGA-MPs (2.5 µg), free IAV PP (2.5 µg), and SEB (0.5 µg) as a control. CXCR5, CCR7, and CXCR3 antibodies were included during stimulation. After 18 h, cells were antibody-stained (Table S6), fixed with 1% paraformaldehyde, acquired in a BD Symphony S6 flow cytometer, and analyzed using FlowJo V10. PLGA without peptides and unstimulated controls set the gate for cytokine-expressing T cells. The antibody panel used for the AIM assay is listed in table S5.

### CFSE T cell proliferation assay

Splenocytes and lung cells were labelled with 0.5 µM CFSE in PBS for 5 mins at room temperature. After washing with RPMI medium, cells were resuspended at 105 cells/mL. CFSE-labeled cells were placed in 96-well plates (200 µl/well) and exposed to IAV PP (10 µg) or PLGA particles (10 µg) for 5 days at 37°C in 5% CO2; RPMI medium was added to control cells. Cells were then stained with APC-conjugated CD4+ antibody, BV780-conjugated CD8α antibody, and Fixable Viability Dye at 4°C for 30 minutes before flow cytometry analysis. Proliferating CD4+ T or CD8+ T cells were assessed by CFSE fluorescence decrease (*70*).

### Quantitative real-time PCR of IFNγ, TNFα, and viral RNA

Mice tissues were homogenized for RNA extraction using TRIzol reagent (Thermo Fisher, 15596018). RNA was reverse transcribed to cDNA using Prime Script^TM^ RT Reagent Kit with gDNA Eraser (Perfect Real Time-RR047A, Takara-Bio) and diluted 5-fold with nuclease-free water (MP Biomedicals, 112450204). Gene expression was studied using PowerUp^TM^ SYBR Green Master Mix (Applied Biosystems^TM^, A25742) with 18s rRNA as an internal control.

### Free IAV PP or PLGA-loaded PP stimulation of spleen and lung cells from mice

Frozen splenocytes and lung cells were resuspended in RPMI 1640 medium with 10% FBS and antibiotics (penicillin and streptomycin) (Thermo Fisher Scientific, 15140122). Cells were counted using a hemocytometer and diluted to 10^7^/mL. Round-bottomed microtiter plate wells (Nunc, 168136) were added with 100 µl cells and stimulated with 100 µl Free T cell peptides (10 µg/mL), PLGA carrying 0.625-10 µg peptide, and Blank PLGA. Antigen stimulation was performed in duplicate wells and incubated in 5% CO2 for 6 h at 37 °C.

### Plaque Assay

Supernatants were diluted tenfold in Opti-MEM and used to infect MDCK cells in 12-well plates for 1 hr at 37°C. The virus inoculum was removed, and cells were covered with 1 mL MEM (Gibco, 61100053) containing 0.6% Oxoid agar (Thermo Scientific, LP0028), 1 μg/mL TPCK trypsin, 0.01% DEAE-Dextran hydrochloride (Sigma Aldrich, D9885), and 0.5% NaHCO3 (MP Biomedicals, 194553). After 48 hrs, cells were fixed with 4% PFA and stained with crystal violet.

### Histopathology

Lung samples were fixed in 10% buffered PFA, embedded in paraffin, and sectioned to 3 μm using a microtome. Sections were stained with Haematoxylin and Eosin and examined microscopically (*59*). Clinical scoring for mouse lungs was based on vascular infiltration, alveolar infiltration, and interstitial pneumonia, rated 1-4 (1 = mild to 4 = very severe). A blinded veterinary pathologist performed analysis.

### Statistical analysis

In this study, all statistical analyses were performed using either Prism 9 (GraphPad Software) or R statistical software (R Foundation for Statistical Computing). Multiple comparisons of three or more groups were performed using the one-way or two-way ANOVA test with Tukey’s multiple comparisons test or Šídák’s multiple comparisons test, and comparisons of survival curves were performed using log-rank (Mantel-Cox) tests, as indicated in each figure legend. All tests were performed at an alpha level of 0.05.

## Author contributions

R.T.Y., R.A., and S.T. conceptualized the study. R.T.Y. performed most of the experiments and interpreted the data. M.S., S.K.N., R.N., and A.K. assisted with mouse experiments, and U.S. performed T cell activation assays using human samples. R.C. and A.B.R. assisted in the LCMS experiment and analysis. R.R. cultured the HK/19 (H3N2 virus) for mouse experiments, and R.T.Y. performed the statistical analysis. R.T.Y. and S.T. cowrote the manuscript. S.T. supervised the project and acquired funding. All authors contributed to the review of the manuscript.

## Supplemental Information

Document S1. Figures S1-S12 and Tables 1-5

## Data and materials availability

All data associated with this study are present in the paper or the supplementary materials. The T cell prediction data, LCMS analysis data, and the R scripts used for this study will be provided upon request.

## Disclaimer

The findings and conclusions in this report are those of the authors and do not necessarily represent the views of the Centers for Disease Control and Prevention.

