## Supplementary figures and tables for "Broad heterologous protection against Influenza A viruses by an adjuvant-free modular mucosal T-cell vaccine platform"

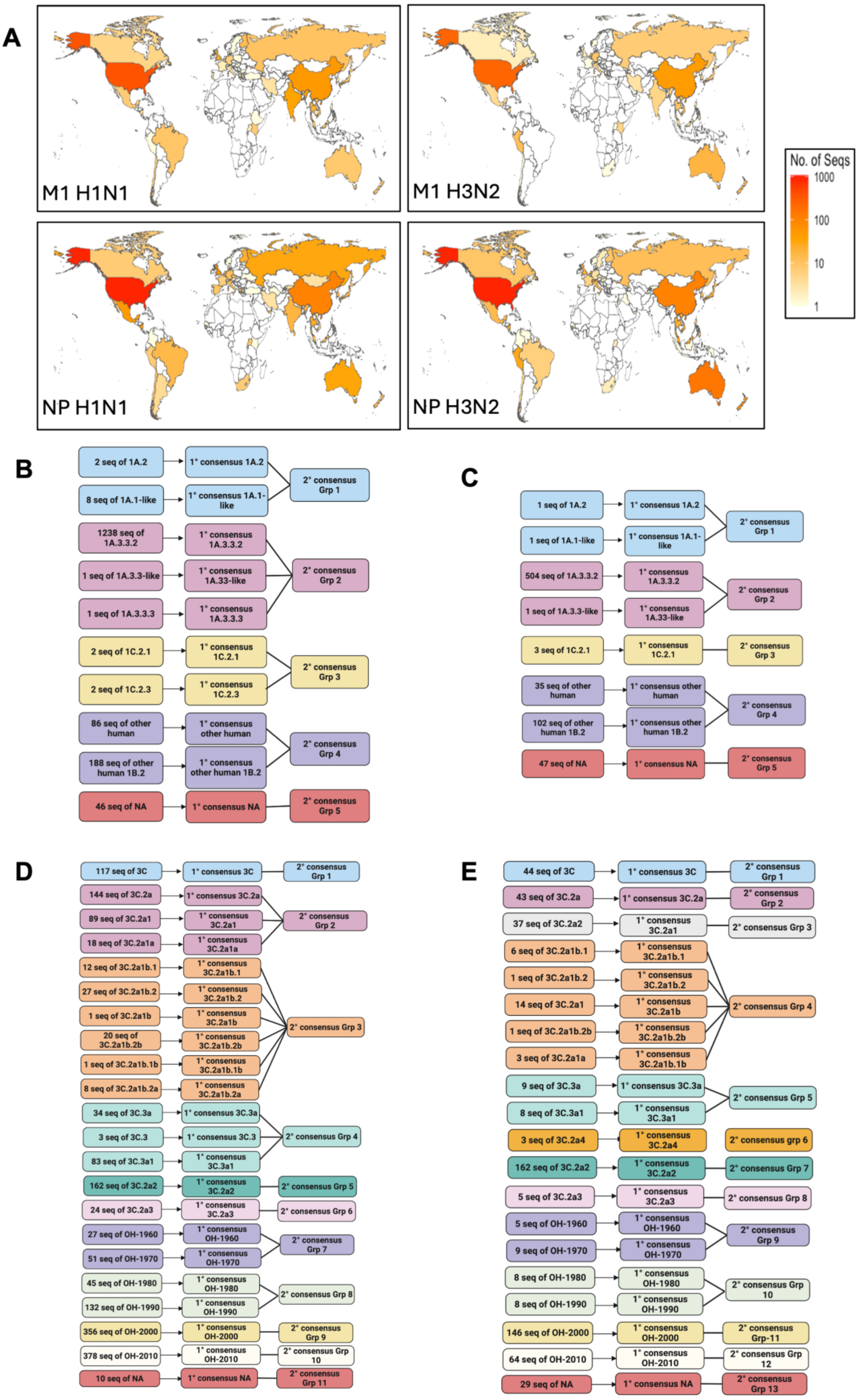

**Figure S1. Geographical Mapping and MMCS-Based Consensus Generation of NP/M1 Proteins in H1/H3 human IAV: (A)** Global distribution heatmaps of influenza virus isolates for H1N1 and H3N2 virus strains (NP and M1). The maps depict the number of sequences reported per country, with a gradient from white (low) to red (high) intensity indicating relative abundance. **(B-E)** Micro consensus sequence generation step followed for **(B)** NP protein of H1N1, **(C)** M1 protein of H3N2, **(D)** NP Protein of H1N1, and **(E)** M1 protein of H3N2.

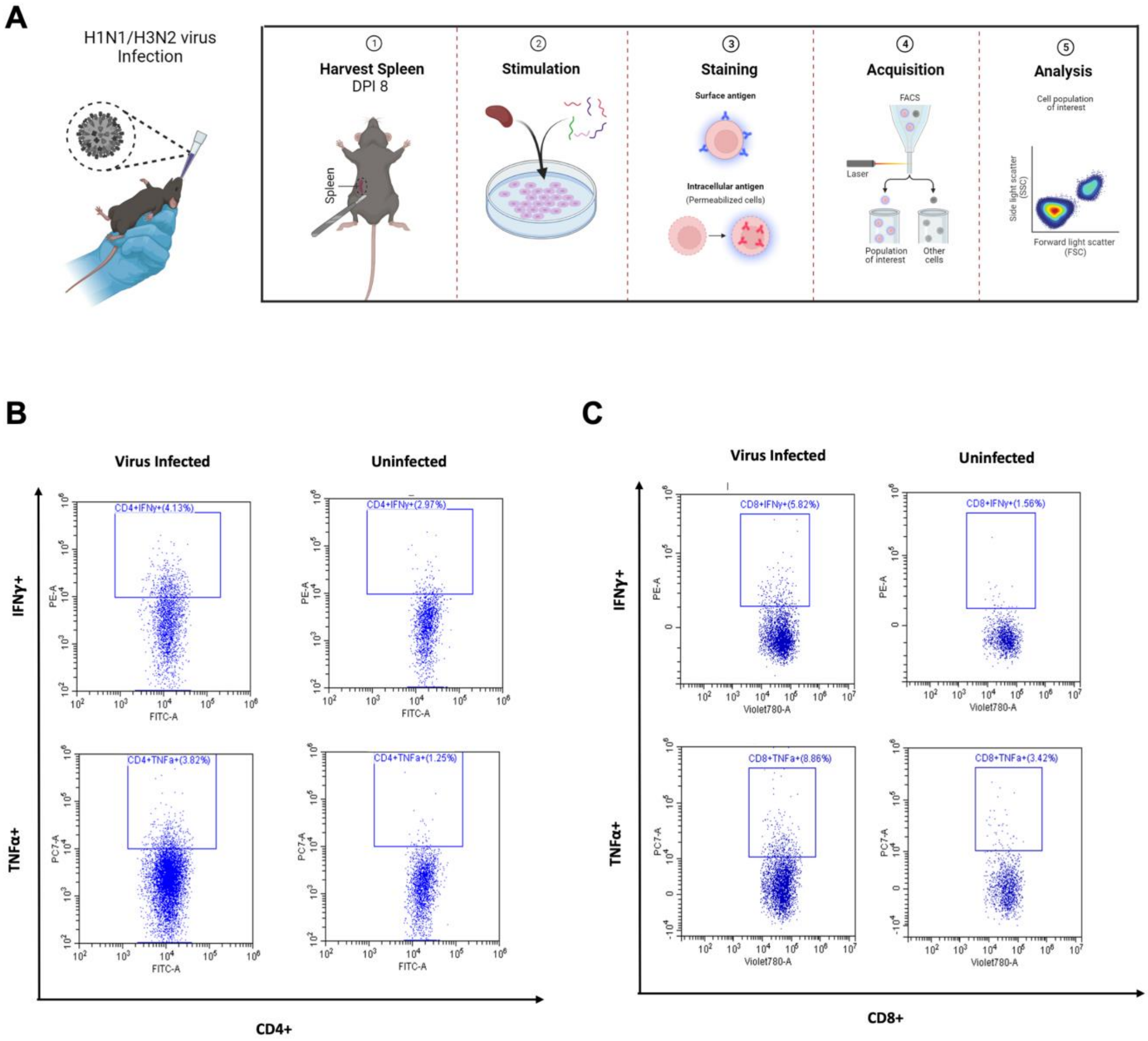

**Figure S2. Experimental outline for validation of CD4+ and CD8+ bispecific peptides: (A)** A schematic illustrates the workflow for evaluating T cell responses following H1N1 virus infection in mice. **(B-C)** Bottom panels show representative intracellular cytokine staining (IFN-γ and TNF-α) in virus-infected versus uninfected mice for both CD4+ (B) and CD8+ T (C) cells, highlighting NP and M1 antigen-specific responses.

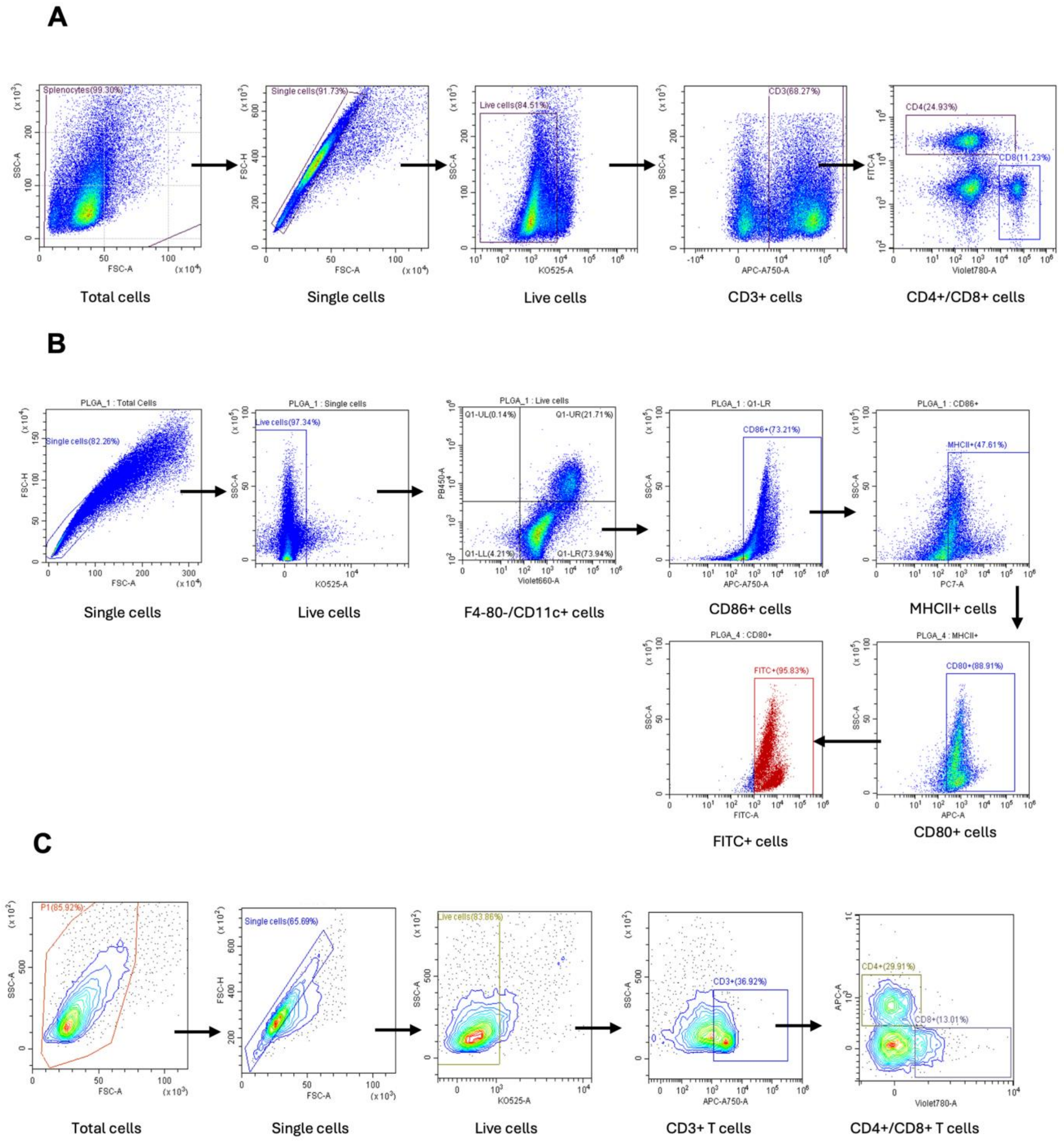

**Figure S3: Gating strategy used for Flow cytometry**: (A) Flow cytometry plots show the sequential gating strategy used to identify CD4+ and CD8+ T cell populations from splenocytes of infected and uninfected mice. Initial gating includes lymphocyte gating, single-cell distribution, and live-cell selection, followed by gating on CD3+ T cells. Subsequent panels show CD4+ and CD8+ T cells. (B) Gating strategy followed for the quantifying the mature BMDCs in coculture experiments, we first looked for F4-80-/CD11c+ population followed for 3 activation markers such as CD80+CD86+MHCII+ in activated BMDC cells. (**C)** Gating strategy used to identify proliferating CD4+ and CD8+ T cells. Total cells were gated for singlets, live cells, CD3+ T cells, and subsequently CD4+/CD8+ populations.

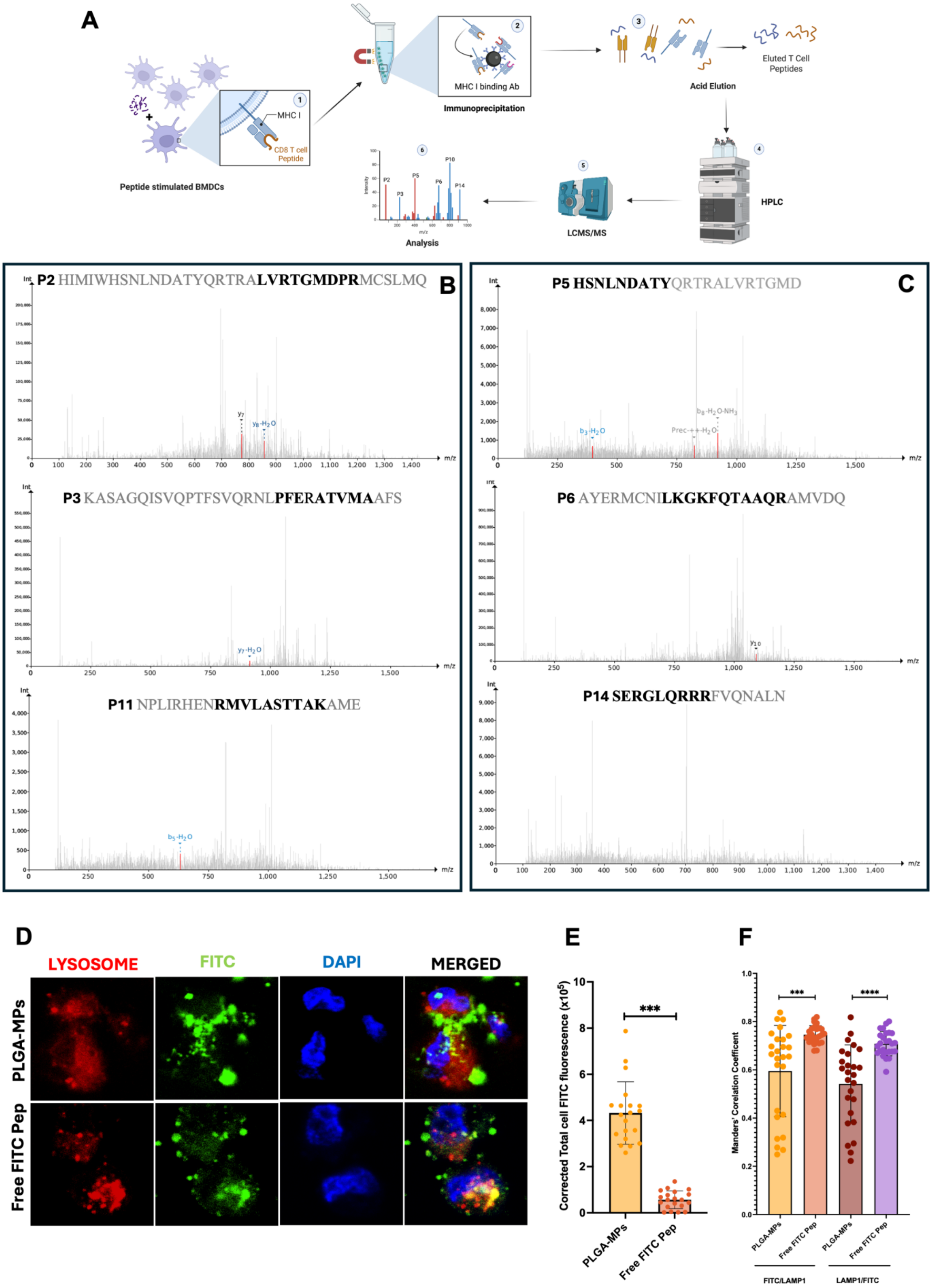

A
Peptide stimulated BMDCs
MHC I
CD8 T cell Peptide
MHC I binding Ab
Immunoprecipitation
Acid Elution
Eluted T Cell Peptides
HPLC
LCMS/MS
Analysis
B
P2 HIMIWHSNLNDATYQRTRALVRTGMDPRMCSLMQ
P3 KASAGQISVQPTFSVQRNLPFERATVMAAFS
P11 NPLIRHENRMVLASTTAKAME
C
P5 HSNLNDATYQRTRALVRTGMD
P6 AYERMCNILKGKFQTAAQRAMVDQ
P14 SERGLQRRRFVQNALN
Int
m/z
D
LYSOSOME
FITC
DAPI
MERGED
PLGA-MPs
Free FITC Pep
E
Corrected Total cell FITC fluorescence (x10^5)
***
PLGA-MPs
Free FITC Pep
F
Manders' Corelation Coefficent
***
****
PLGA-MPs
Free FITC Pep
FITC/LAMP1
LAMP1/FITC

**Figure S4**: **PLGA-mediated delivery enhances antigen processing, cross-presentation, and endosomal escape (A)** Schematic representation of the immunopeptidome assay. **(B, C)** LC-MS spectra showing detection of known influenza-derived CD8+ T cell epitopes (bold) in PLGA-MPs encapsulated IAV PP. Identified peptides correspond to selected regions of the NP and M1 proteins (P2, P3, P11 for H1N1; P5, P6, P14 for H3N2), confirming successful processing and presentation on MHC-I molecules. Annotated peaks indicate experimentally validated epitope sequences. **(D)** Confocal microscopy images showing intracellular localization of FITC-labelled peptide delivered either as free peptide or encapsulated within PLGA-MPs in THP-1 cells. Lysosomes are stained in red using LAMP-1, peptide signal in green (FITC), and nuclei in blue (DAPI). Merged images demonstrate reduced colocalization of PLGA-delivered peptide with lysosomes, indicating enhanced endosomal escape compared to free peptide. **(E)** Quantification of cellular uptake based on corrected total cell FITC fluorescence = Integrated density – (Area of selected cell x Mean background fluorescence). PLGA-MP encapsulated peptides show significantly higher intracellular FITC signal compared to free peptide, indicating enhanced uptake efficiency. **(F)** Quantification of colocalization using Manders' correlation coefficients. FITC/LAMP1 denotes the fraction of FITC (green) signal overlapping with LAMP-1 (red), and LAMP1/FITC denotes the fraction of LAMP-1 (red) signal overlapping with FITC (green). PLGA-MP–delivered peptide shows significantly reduced colocalization compared to free peptide, indicating enhanced endosomal escape and cytosolic availability. Quantification was performed across *25* independent cells from 6-7 images. Data are presented as mean ± SEM; statistical significance determined by one-way ANOVA with Tukey's multiple comparison test (**$p < 0.001$).

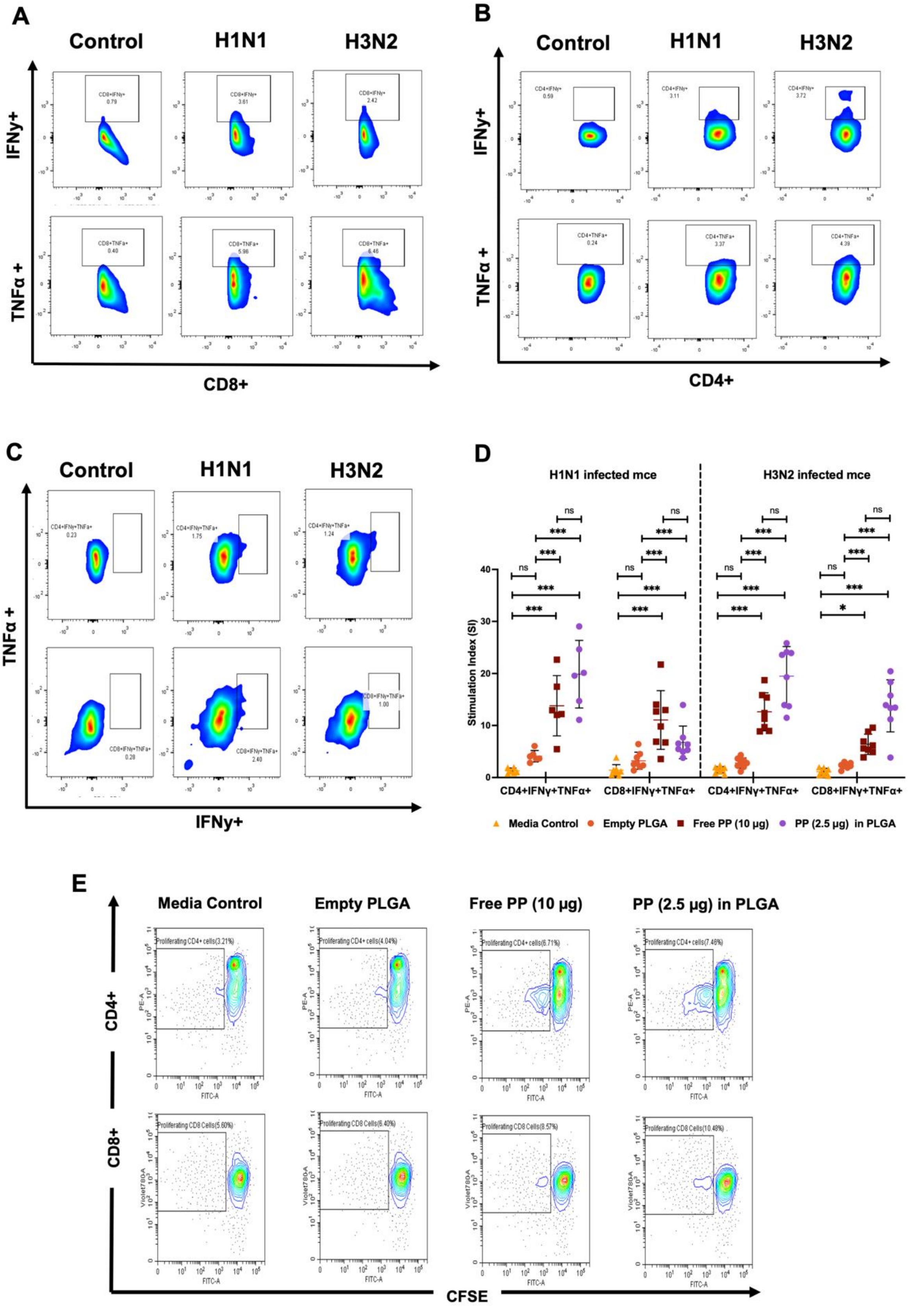
A
Control
H1N1
H3N2
IFNy+
TNFα +
CD8+
CD8+IFNy+
0.79
3.61
2.42
CD8+TNFa+
0.40
5.96
6.46
B
Control
H1N1
H3N2
IFNy+
TNFα +
CD4+
CD4+IFNy+
0.59
3.11
3.72
CD4+TNFa+
0.24
3.37
4.39
C
Control
H1N1
H3N2
TNFα +
IFNy+
CD4+IFNy+TNFa+
0.23
1.75
1.24
CD8+IFNy+TNFa+
0.28
2.40
1.00
D
H1N1 infected mce
H3N2 infected mce
Stimulation Index (SI)
0
10
20
30
40
ns
***
*
CD4+IFNγ+TNFα+
CD8+IFNγ+TNFα+
Media Control
Empty PLGA
Free PP (10 μg)
PP (2.5 μg) in PLGA
E
Media Control
Empty PLGA
Free PP (10 μg)
PP (2.5 μg) in PLGA
CD4+
CD8+
CFSE
Proliferating CD4+ cells(3.21%)
Proliferating CD4+ cells(4.04%)
Proliferating CD4+ cells(6.71%)
Proliferating CD4+ cells(7.46%)
Proliferating CD8 Cells(5.60%)
Proliferating CD8 Cells(6.40%)
Proliferating CD8 Cells(8.57%)
Proliferating CD8 Cells(10.48%)
PE-A
Violet780A
FITC-A

**Figure S5: Functional characterization and proliferation of CD4+ and CD8+ T cells in response to ma/Cal/09 H1N1 and HK/19 H3N2 influenza virus infection: (A, B)** Representative intracellular cytokine staining (ICS) dot plots showing IFN-γ and TNF-α expression in $CD4^+$ and $CD8^+$ T cells from control, H1N1-infected, and H3N2-infected mice, indicating virus NP and M1 protein-specific T cell activation. **(C, D)** Representative dot plot and quantification of polyfunctional T cells expressing IFN-γ and TNF-α in $CD4^+$ and $CD8^+$ populations from H1N1- and H3N2-infected mice. Comparisons were made between media control, free peptide (PP), peptide-loaded PLGA-MPs, and empty PLGA conditions. Data are presented as fold increases relative to the control. Statistical significance is indicated (*$p<0.05$, **$p<0.01$, ***$p<0.001$). **(E)** Representative contour plots showing T cell proliferation in response to PP-loaded PLGA-MPs, free PP, media control, and empty PLGA. Increased CFSE dilution indicates greater proliferation of $CD4^+$ and $CD8^+$ T cells in response to antigenic stimulation. There were 8 mice per group for all the above studies.

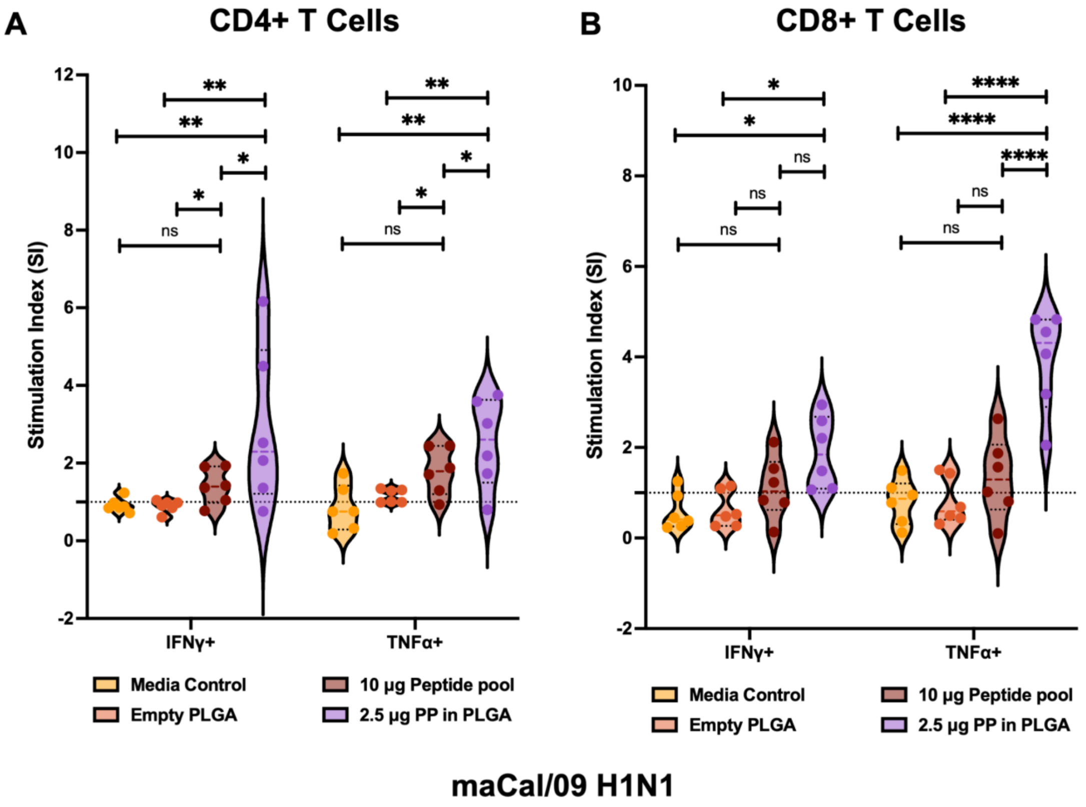

**Figure S6: Comparison of T cell activation between Free IAV PP and IAV PP Loaded in PLGA-MPs in lung cells post maCal/09 H1N1 infection:** CD4+ T cell activation **(A)** and CD8+ T cell activation **(B)** in lung cells from maCal/09 H1N1 infected mice was measured by intracellular cytokine staining. Lungs were stimulated with IAV PP in different formulations, and cytokine production (IFN-γ+ and TNF-α+) was measured by flow cytometry.

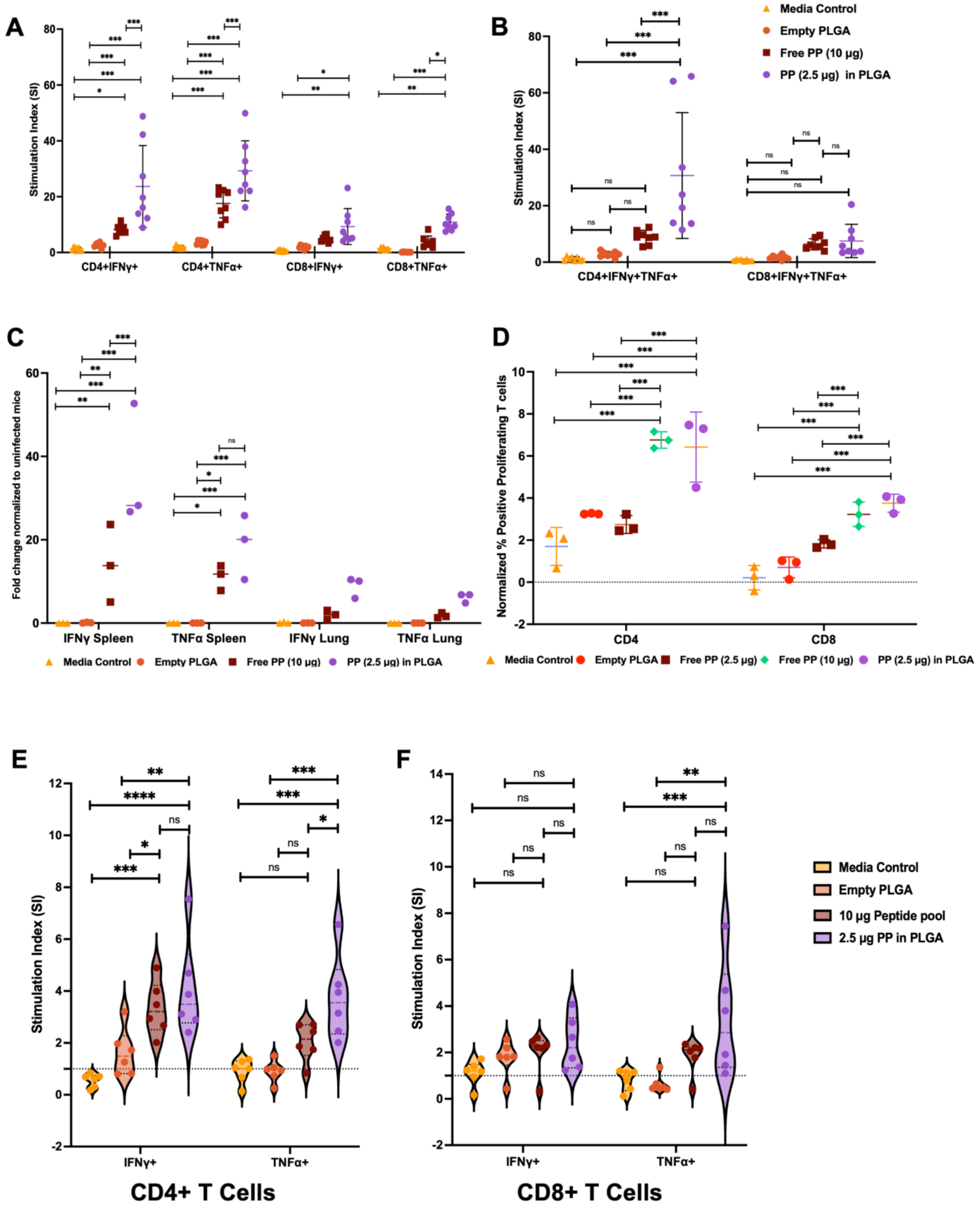

**Figure S7: Comparison of T cell activation and proliferation induced by the free IAV PP and IAV PP loaded in PLGA-MPs in splenocytes from the X31 virus-infected mice:** **(A)** ICS staining of CD4 and CD8 T cells secreting IFNγ+/TNFα+ cytokine following *ex vivo* peptide stimulation in mice splenocytes 8DPI with X31 virus. **(B)** Polyfunctional CD4/CD8 T cells activation post peptide stimulation **(C)** qRT-PCR quantification of IFN-γ and TNF-α mRNA expression. **(D)** CFSE-based T cell proliferation assay. **(E)** CD4+ T cell activation and **(F)** CD8+ T cell activation in lung cells from X31/PR8-infected mice were measured by intracellular cytokine staining. Data are presented as mean ± SEM. An ordinary two-way ANOVA test determined statistical significance. $^*p < 0.05$, $^{**}p < 0.01$, $^{***}p < 0.001$.

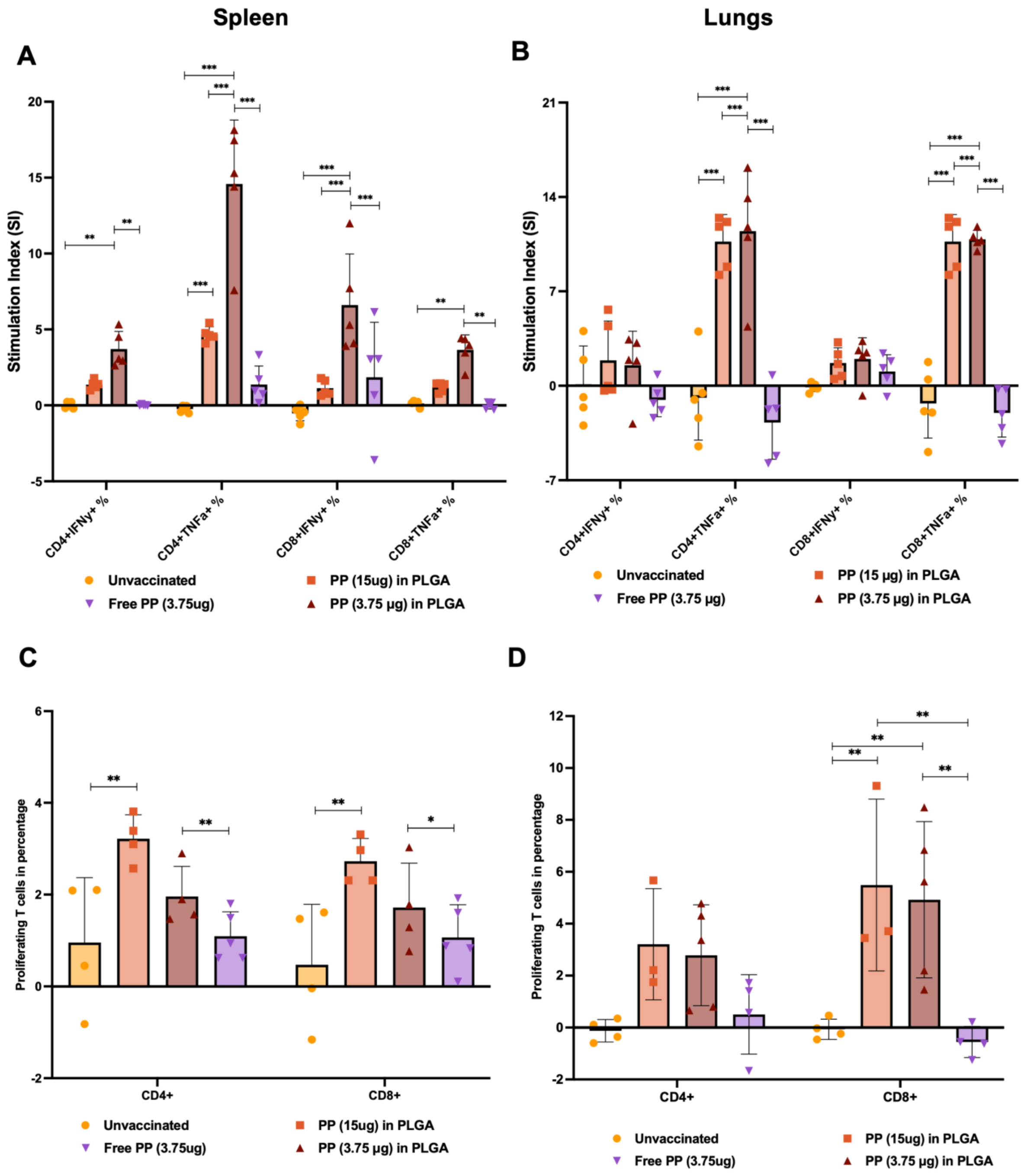

**Figure S8: T cell cytokine responses and proliferation following vaccination with PLGA-encapsulated IAV peptide pool.** (A, B) Frequency of cytokine-producing CD4$^+$ and CD8$^+$ T cells following ex vivo co-culture of CD3+ enriched T cells from splenocytes **(A)** and lungs **(B)** from vaccinated mice with naïve bone marrow-derived dendritic cells (BMDCs) pulsed with influenza A virus peptide pool (IAV PP). CD3+ T cells were isolated from mice vaccinated with 15 µg PP in PLGA, 3.75 µg PP in PLGA, and 3.75 µg free PP. Bar graphs represent the percentage of single-positive IFN-γ+ and TNF-α+ CD4+ and CD8+ T cells. **(C, D)** T cell proliferation analysis by CFSE dilution assay following ex vivo IAV PP stimulation of splenocytes from vaccinated or unvaccinated mice. Frequencies of proliferating (CFSE) CD4+ and CD8+ T cells in splenocytes **(C)** and lungs **(D)** are shown. Data represent mean ± SEM. Statistical significance was calculated using Two-way ANOVA **(A, B)** and one-way ANOVA with post-hoc test **(C, D)**. $^*p < 0.05$, $^{**}p < 0.01$, $^{***}p < 0.001$.

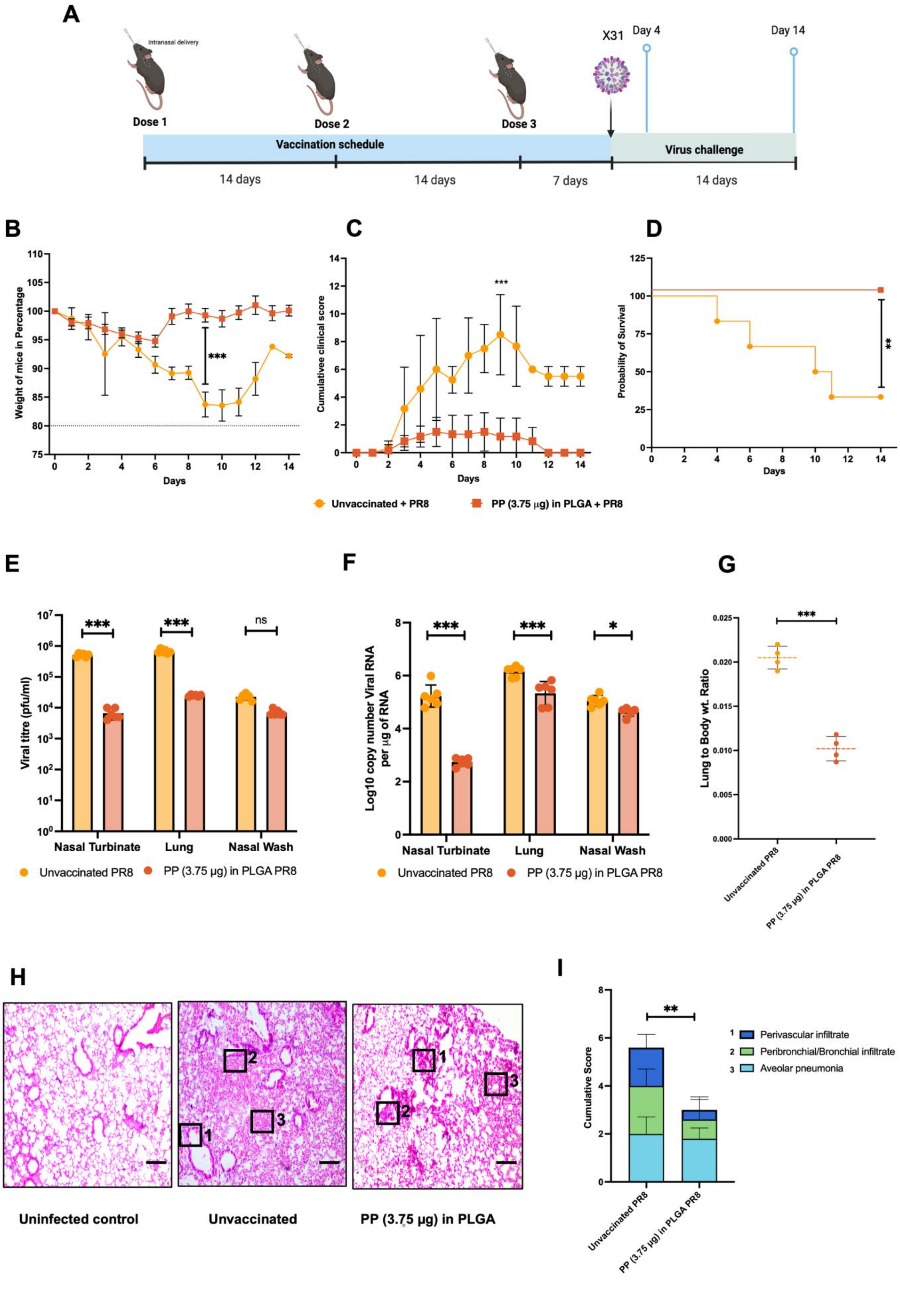

A
Intranasal delivery
Dose 1
Dose 2
Dose 3
X31
Day 4
Day 14
Vaccination schedule
Virus challenge
14 days
14 days
7 days
14 days
B
Weight of mice in Percentage
Days
***
C
Cumulativee clinical score
Days
***
D
Probability of Survival
Days
**
Unvaccinated + PR8
PP (3.75 µg) in PLGA + PR8
E
Viral titre (pfu/ml)
Nasal Turbinate
Lung
Nasal Wash
***
***
ns
Unvaccinated PR8
PP (3.75 µg) in PLGA PR8
F
Log10 copy number Viral RNA per µg of RNA
Nasal Turbinate
Lung
Nasal Wash
***
***
*
Unvaccinated PR8
PP (3.75 µg) in PLGA PR8
G
Lung to Body wt. Ratio
***
Unvaccinated PR8
PP (3.75 µg) in PLGA PR8
H
Uninfected control
Unvaccinated
PP (3.75 µg) in PLGA
I
Cumulative Score
**
Perivascular infiltrate
Peribronchial/Bronchial infiltrate
Aveolar pneumonia
Unvaccinated PR8
PP (3.75 µg) in PLGA PR8

**Figure S9: Evaluation of Immune Protection induced by intranasal vaccination against PR8 viruses**: **(A)** Schematic of the vaccination and PR8 IAV virus challenge study in mice. Mice received 3 intranasal vaccine doses at 14-day intervals. **(B)** Percentage weight change in mice over 14 days post-virus challenge. Weight loss trends were compared between unvaccinated and vaccinated groups. **(C)** Cumulative clinical scores over 14 days post-challenge, evaluating symptoms among groups receiving different vaccine formulations and the unvaccinated group. **(D)** Survival curve of mice over 14 days post-virus challenge. **(E)** Viral titers pfu/mL and Viral RNA copy numbers **(F)** in the nasal turbinate, lung and nasal wash indicate viral load reduction and reflect vaccine efficacy in controlling viral replication. **(G)** Lung-to-body weight ratios, indicating pulmonary inflammation severity among groups. **(H)** H and E staining images **(I).** Cumulative histopathological scoring of lung tissues for perivascular infiltrate, peribronchial/bronchiolar infiltrate, and alveolar pneumonia severity. Data are presented as mean ± SEM. Statistical significance was determined by an ordinary two-way ANOVA test. *p < 0.05, **p < 0.01, ***p < 0.001 and Scale bar = 200 µM

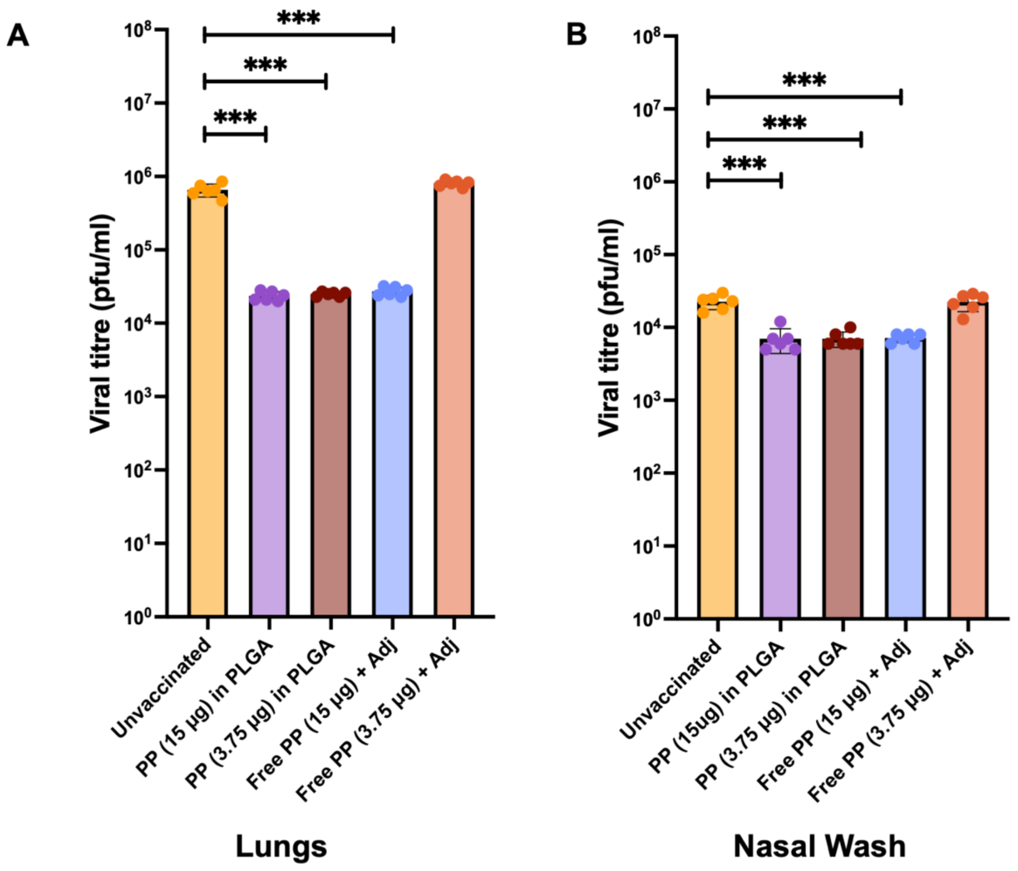

**Figure S10: Quantification of infectious virus in lung and nasal wash samples following maCal/09 H1N1 influenza virus challenge.** Viral titers in the lungs **(A)** and nasal wash **(B)** were measured by plaque assay. Data are presented as $\log_{10}$ PFU/mL. Mice were treated with various formulations: free PP (15 or 3.75 µg), or PP-loaded PLGA-MPs (3.75 µg PP in PLGA). n=6 mice per group, Bars represent mean ± SEM. Statistical comparisons were performed using one-way ANOVA with significance indicated as *p < 0.05, **p < 0.01, ***p < 0.001.

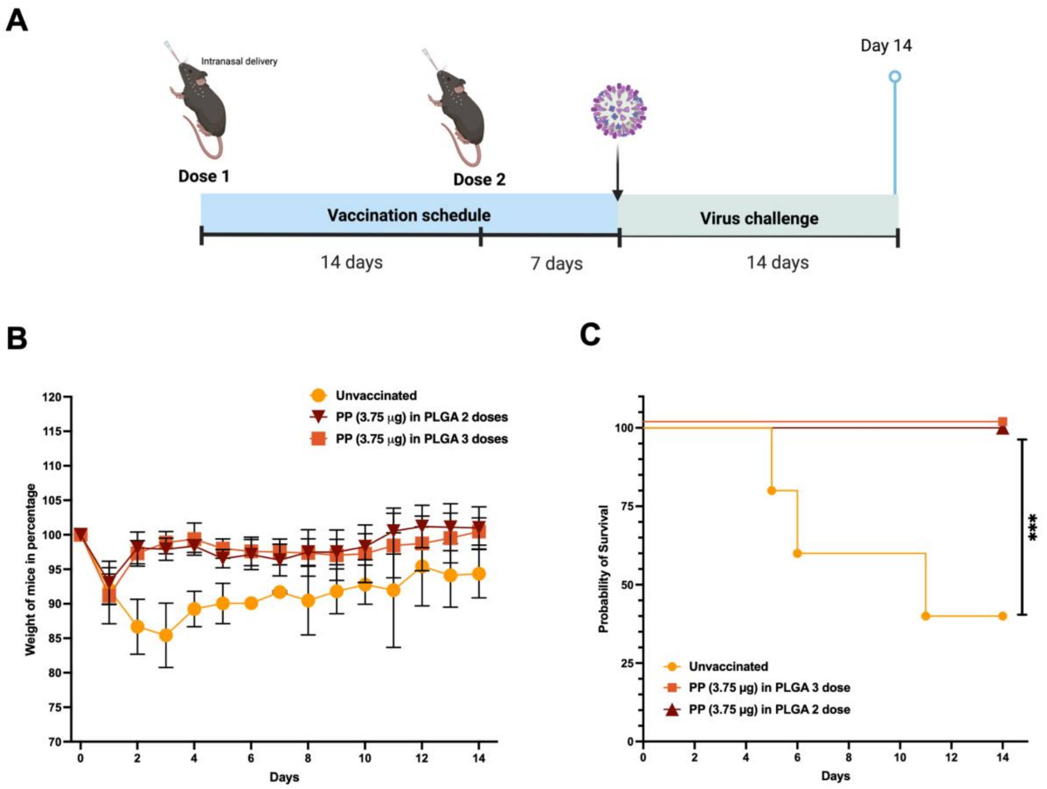

**Figure S11: Efficacy of reduced vaccine dose utilizing PLGA-MPs: (A)** Schematic representation of the vaccination and challenge timeline. Mice received two intranasal doses of peptide loaded PLGA nanoparticles (PP in PLGA) on Days 0 and 14. On Day 21, mice were challenged with influenza virus and monitored for 14 days post-infection. **(B)** Percent body weight change following viral challenge. Mice vaccinated with PP (3.75 µg) in PLGA were compared to unvaccinated controls. Body weight was monitored daily and plotted as percentage relative to initial weight. Data represent mean ± SEM (n = 5–6 mice per group). **(C)** Kaplan-Meier survival curve showing the percentage of surviving mice post-influenza challenge. Vaccinated groups demonstrated enhanced protection compared to unvaccinated controls.

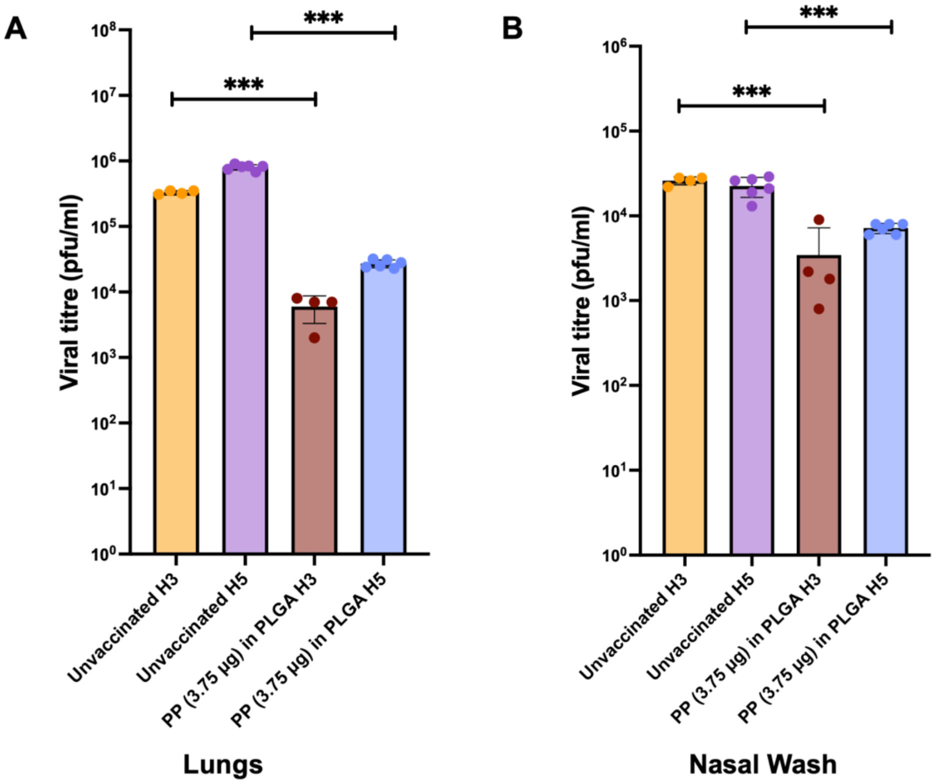

**Figure S12: Virus titer quantification post H3 and H5 virus challenge. (A, B)** Viral titers in the lungs **(A)** and nasal washes **(B)** were determined by plaque assay and presented as $\log_{10}$ pfu/mL. Mice were unvaccinated or PP-loaded PLGA-MPs (3.75 µg PP in PLGA). n= 4 or 6 mice per group, and the bars indicate mean ± SEM. Statistical analysis was conducted using one-way ANOVA, with significance levels denoted as $^*p < 0.05$, $^{**}p < 0.01$, $^{***}p < 0.001$.

### Supplementary Tables:

| Sr. No | MHC I | MHC II |
|---|---|---|
| 1 | HLA-A*01:01 | HLA-DRB1*01:01 |
| 2 | HLA-A*02:01 | HLA-DRB1*03:01 |
| 3 | HLA-A*02:03 | HLA-DRB1*04:01 |
| 4 | HLA-A*02:06 | HLA-DRB1*04:05 |
| 5 | HLA-A*03:01 | HLA-DRB1*07:01 |
| 6 | HLA-A*11:01 | HLA-DRB1*08:02 |
| 7 | HLA-A*23:01 | HLA-DRB1*09:01 |
| 8 | HLA-A*24:02 | HLA-DRB1*11:01 |
| 9 | HLA-A*26:01 | HLA-DRB1*12:01 |
| 10 | HLA-A*30:01 | HLA-DRB1*13:02 |
| 11 | HLA-A*30:02 | HLA-DRB1*15:01 |
| 12 | HLA-A*31:01 | HLA-DRB3*01:01 |
| 13 | HLA-A*32:01 | HLA-DRB3*02:02 |
| 14 | HLA-A*33:01 | HLA-DRB4*01:01 |
| 15 | HLA-A*68:01 | HLA-DRB5*01:01 |
| 16 | HLA-A*68:02 | HLA-DQA1*05:01/DQB1*02:01 |
| 17 | HLA-B*07:02 | HLA-DQA1*05:01/DQB1*03:01 |
| 18 | HLA-B*08:01 | HLA-DQA1*03:01/DQB1*03:02 |
| 19 | HLA-B*15:01 | HLA-DQA1*04:01/DQB1*04:02 |
| 20 | HLA-B*35:01 | HLA-DQA1*01:01/DQB1*05:01 |
| 21 | HLA-B*40:01 | HLA-DQA1*01:02/DQB1*06:02 |
| 22 | HLA-B*44:02 | HLA-DPA1*02:01/DPB1*01:01 |
| 23 | HLA-B*44:03 | HLA-DPA1*01:03/DPB1*02:01 |
| 24 | HLA-B*51:01 | HLA-DPA1*01:03/DPB1*04:01 |
| 25 | HLA-B*53:01 | HLA-DPA1*03:01/DPB1*04:02 |
| 26 | HLA-B*57:01 | HLA-DPA1*02:01/DPB1*05:01 |
| 27 | HLA-B*58:01 | HLA-DPA1*02:01/DPB1*14:01 |

**Table S1:** HLA MHC I/II Panel used for the T cell epitope prediction.

| No. | Cross protection against Influenza subtypes |
|---|---|
| P1 | H2N2, H9N1, H3N2 , H3N1, H1N1, H7N2, H5N1, H9N1, H3N1, H2N3 |
| **P2** | H1N1, H3N2, H5N1, H6N1, H9N2, |
| **P3** | H1N1, H1N2, H3N2, H6N1, H5N1, H9N2 |
| P4 | H1N1, H2N3, H7N9, H9N1, H3N1, H3N2, H2N2, |
| **P5** | H5N1, H1N1, H3N1, H3N2, H5N1 H6N1 H9N2, H9N1 |
| **P6** | H3N2, H1N1, H3N1, H5N1 |
| P7 | H5N1 H3N2 H6N1 H9N2 |
| P8 | H1N1, H1N2, H2N2 H3N1, H3N2 H9N1 H5N1 H1N2v, H3N8 |
| P9 | H1N1, H1N2, H2N2, H3N1, H3N2, H5N1, H9N1, |
| P10 | H1N1 H1N2, H3N1 H3N2 H9N2, H3N8, H4N6, H5N1, H6N2 |
| **P11** | H3N8, H3N2, H9N2, H4N6, H5N1, H6N2, H1N1, H1N2, H3N1, H2N2,H9N1, H2N2, H9N1, |
| P12 | H1N1, H1N2, H1N2v, H3N1, H3N2, H9N2, , H2N3, H9N1, H2N2, |
| P13 | H2N2, H3N1, H3N2, H5N1, H6N9, H1N1, H9N1, H1N2, H3N8, H4N6, H9N1 |
| **P14** | H2N3, H3N2, H1N1, H3N2, H1N2, H2N2 H3N1, H5N1, H9N1 H6N2, H4N6, H3N8 |
| P15 | H1N1, H1N2, H2N2, H9N1, H5N1,H3N2, H3N1, H2N3 |
| P16 | H1N1, H1N2, H3N1, H3N2, H9N2, H2N2, H5N1, H9N1, |

**Table S2:** The selected 16 bispecific T cell peptides have shown to be present in multiple Influenza subtypes. The selected 6 peptides for the final vaccine formulation is highlighted in bold.

| No. | Protein | Influenza type | Overlapping Epitope | Size | MHC I Allele | CD4+ epitope Sequence | MHC II Allele | CD8+ epitope Sequence |
|---|---|---|---|---|---|---|---|---|
| P1 | NP protein | H1N1 | MIGGIGRFYIQMCTELKLSDYEGRLI | 26 | HLA-A*0101 | CTELKLSDY | HLA-DRB1*04:01 | GRFYIQMCTELKLSD |
| | | | | | HLA-A*2402 | FYIQMCTEL | | |
| **P2** | | | HIMIWHSNLNDATYQRTRALVRTGMDPRMCSLMQ | 34 | HLA-A*02:01 | GMDPRMCSL | HLA-DRB1*02:02 | MIWHSNLNDATYQRT |
| | | | | | | | HLA-DRB5*01:01 | DATYQRTRALVRTGM |
| **P3** | | | KASAGQISVQPTFSVQRNLPFERATVMAAFS | 31 | HLA-A*26:01 | ERATVMAAF | HLA-DRB1*01:01 | ASAGQISVQPTFSVQRNLPF |
| | | | | | HLA-B*07:02 | LPFERATVM | | |
| P4 | | H3N2 | GIGRFYIQMCTELKLSDHEGR | 21 | HLA-A*24:02 | FYIQMCTEL | HLA-DRB1*04:01 | GRFYIQMCTELKLSD |
| **P5** | | | HSNLNDATYQRTRALVRTGMD | 21 | HLA-A*01:01 | HSNLNDATY | HLA-DRB5*01:01 | ATYQRTRALVRTG |
| | | | | | HLA-B*07:02 | ATYQRTRAL | HLA-DRB1*07:01 | DATYQRTRALVRTGMD |
| **P6** | | | AYERMCNILKGKFQTAAQRAMVDQ | 24 | HLA-A*39:01 | FQTAAQRAM | HLA-DRB1*13:02 | RMCNILKGKFQTAAQRAM |
| | | | | | HLA-B*0801 | ILKGKFQTA | | |
| p7 | | | IRPNENPAHKSQLVWMACHSAAFEDLRL | 28 | HLA-B*1501 | WMACHSAAF | HLA-DRB1*04:05 | NPAHKSQLVWMACHSAAFED |
| P8 | M1 Protein | H1N1 | TYVLSIIPSGPLKAEIAQRLEDVFAGKNTDL | 31 | HLA-A*11:01 | SIIPSGPLK | HLA-DRB1*08:02 | SGPLKAEIAQRLEDV |
| | | | | | | | HLA-DRB1*01:01 | TYVLSIIPSGPLKAEI |
| P9 | | | LTKGILGFVFTLTVPSERGLQRR | 23 | HLA-A*0201 | GILGFVFTL | HLA-DRB1*04:01 | KGILGFVFTLTVPSER |
| P10 | | | RAVKLYKKLKREITFHGAKEVSLSYS | 26 | HLA-A*2402 | TFHGAKEVS | HLA-DRB1*11:01 | AVYKKLKREITFHGA |
| **P11** | | | NPLIRHENRMVLASTTAKAME | 21 | HLA-A*0301 | MVLASTTAK | HLA-DRB1*03:01 | IRHENRMVLASTTAKA |
| | | | | | HLA-B*3901 | IRHENRMVL | | |
| P12 | | | GTHPSSSAGLKDDLLENLQAYQKRMGVQMQR | 31 | HLA-B*0801 | YQKRMGVQM | HLA-DRB4*01:01 | LQAYQKRMGVQMQRF |
| | | | | | HLA-A*2601 | DLLENLQAY | | |
| P13 | | H3N2 | DLEALMEWLKTRPILSPLTKGILG | 24 | HLA-B*03:01 | ALMEWLKTR | HLA-DRB1*07:01 | MEWLKTRPILSPL |
| | | | | | HLA-A*40:01 | MEWLKTRPI | | |
| | | | | | HLA-B*0702 | KTRPILSPL | | |
| **P14** | | | SERGLQRRRFVQNALN | 16 | HLA-B*2705 | RRRFVQNAL | HLA-DRB3*02:02 | GLQRRRFVQNALN |
| P15 | | | NPLIKHENRMVLASTTAKAME | 21 | HLA-A*0301 | RLEDVFAGK | HLA-DRB1*13:02 | IKHENRMVLASTTAKAM |
| P16 | | | LLENLQTYQKRMGVQMQRFK | 20 | HLA-B*15:01 | YQKRMGVQM | HLA-DRB5*01:01 | QKRMGVQMQRFK |
| | | | | | HLA-A*2402 | RMGVQMQRF | | |

**Table S3:** Synthesized 16 bispecific T cell peptides used for our ex vivo validation study.

| **ID** | **age** | **Gender** | **race** | **Season** | **HLA-A** | | **HLA-B** | | **HLA-C** | |
|---|---|---|---|---|---|---|---|---|---|---|
| **3430928** | 42 | F | B | 2012 | 02:01 | 02:05 | 07:06 | 18:01 | 05:01 | 07:02 |
| **3430989** | 61 | F | W | 2013 | 02:01 | 68:01:00 | 44:02 (H) | 44:02 (H) | 05:01(H) | 05:01(H) |
| **3430877** | 51 | F | W | 2014 | 02:01 | 24:02:00 | 39:01:00 | 49:01:00 | 07:01 | 12:03 |
| **3431250** | 31 | M | W | 2013 | 02:01 | 03:01 | 07:02 | 15:23 | 07:02 | 07:04 |
| **3430021** | 50 | M | W | 2013 | A*01:01 | A*03:01 | B*07:02(H) | B*07:02(H) | C*07:01 | C*07:02 |

| **Flu Season** | **H1N1** | **H3N2** | **B Lineage** |
|---|---|---|---|
| 2011-12 | A/California/7/2009 (H1N1) pdm09-like | A/Perth/16/2009 (H3N2)-like | B/Brisbane/60/2008-like (Victoria lineage) |
| 2012-13 | A/California/7/2009 (H1N1) pdm09-like | A/Victoria/361/2011 (H3N2)-like | B/Wisconsin/1/2010-like (Yamagata lineage). |
| 2013-14 | A/California/7/2009 (H1N1) pdm09-like | A/Texas/50/2012 (H3N2)-like | B/Massachusetts/2/2012-like (Yamagata lineage). |

| **ID** | **3430928** | **3430989** | **3430877** | **3431250** | **3430021** |
|---|---|---|---|---|---|
| DRB5 | | 1:01 | 2:02 | | *01:01 |
| DQB1 | 2:02 | 3:01 | 3:01 | 3:02 | *02:01 |
| | 3:19 | 6:02 | 5:02 | 6:03 | *06:02 |
| DQA1 | 2:01 | 1:02 | 1:02 | 1:03 | *01:02 |
| | 4:01 | 3:03 | 3:03 | 3:01 | *05:01 |
| DRB1 | 8:04 | 4:01 | 4:07 | 04:04P | *03:01 |
| | 13:03 | 15:01 | 16:01 | 13:01 | *15:01 |
| DRB3 | 2:02 | | | 1:01 | *01:01 |
| | 2:02 | | | | |
| DRB4 | | 1:03 | 1:03 | 1:03 | |
| DPB1 | 1:01 | 4:01 | 4:01 | 02:01P | *04:01 |
| | 2:01 | 10:01 | 35:01:00 | 11:01 | *04:01 |
| DPA1 | 2:02:02 | 1:03 | 1:03 | 1:03 | *01:03(H) |
| | 2:02:04 | 2:01 | 2:01 | 2:01 | *01:03(H) |

**Table S4**: MHC I and II panel of the human cohort selected for our study.

| Laser | Fluorochrome | Marker | Clone | Catlog no. | Company |
|---|---|---|---|---|---|
| R 670/30 | AF647 / APC / NR660 / AF660 / CellTrace Red | CD137 | 4B4-1 | 309810 | Biolegend |
| R 710/50 | AF700 / redFluor710 / R718 / NR710 / APC- R700<br>GhostDye R710 | CD40L | 24-31 | 310846 | Biolegend |
| R 780/60 | APC-Cy7 / APC-Fire750 / APC-H7 / AF750 / APC-AF750<br>GhostDye R780 | L/D | | L34976 | Life Technologies, Thermo Fisher Scientific |
| V 431/28 | BV421 / AF405 / DyLight405 / DAPI | CD69 | FN50 | 310930 | Biolegend |
| V 450/50 | Calcein Violet | | | | |
| V 470/15 | BV480 / Pac Blue / CellTrace Violet | CD4 | RPA-T4 | 569781 | BD |
| V 515/30 | BV510 / GhostDye V510 | | | | |
| V 586/15 | BV570 GhostDye V540 / Zombie Aqua | CD8 | RPA-T8 | 301038 | Biolegend |
| V 780/60 | BV785 / BV786 | | | | |
| UV 379/28 | BUV395 | PDL-1 | 29E.2A3 | 568620 | BD |
| UV 820/60 | BUV805 | CD3 | UCHT1 | 612895 | BD |
| YG 586/15 | PE / BYG584 / NY590 / DL550 / CellTrace Yellow/Calcein Red / Phrodo / Td-Tomato / DsRed | OX-40 | (ACT35) | 350004 | Biolegend |
| YG 780/60 | PE-Cy7 / BYG790 | CD56 | NCAM16.2 | 335791 | BD |

**Table S5:** Antibody panel used for the AIM Assay
